# Engineering multiple species-like genetic incompatibilities in insects

**DOI:** 10.1101/2020.04.03.024588

**Authors:** Maciej Maselko, Nathan Feltman, Ambuj Upadhyay, Amanda Hayward, Siba Das, Nathan Myslicki, Aidan J. Peterson, Michael B. O’Connor, Michael J. Smanski

**Affiliations:** Department of Biochemistry, Molecular Biology, and Biophysics, University of Minnesota, Saint Paul, MN 55108; Biotechnology Institute, University of Minnesota, Saint Paul, MN 55108; Department of Genetics, Cell Biology, and Development, University of Minnesota, Saint Paul, MN 55108

## Abstract

Speciation constrains the flow of genetic information between populations of sexually reproducing organisms. Gaining control over mechanisms of speciation would enable new strategies to manage wild populations of disease vectors, agricultural pests, and invasive species. Additionally, such control would provide safe biocontainment of transgenes and gene drives. Natural speciation can be driven by pre-zygotic barriers that prevent fertilization or by post-zygotic genetic incompatibilities that render the hybrid progeny inviable or sterile. Here we demonstrate a general approach to create engineered genetic incompatibilities (EGIs) in the model insect *Drosophila melanogaster*. Our system couples a dominant lethal transgene with a recessive resistance allele. EGI strains that are homozygous for both elements are fertile and fecund when they mate with similarly engineered strains, but incompatible with wild-type strains that lack resistant alleles. We show that EGI genotypes can be tuned to cause hybrid lethality at different developmental life-stages. Further, we demonstrate that multiple orthogonal EGI strains of *D. melanogaster* can be engineered to be mutually incompatible with wild-type and with each other. Our approach to create EGI organisms is simple, robust, and functional in multiple sexually reproducing organisms.

## Main Text

In genetics, underdominance occurs when a heterozygous genotype (Aa) is less fit than either homozygous genotype (AA and aa). In ‘extreme underdominance’, the heterozygote is inviable while each homozygote has equal fitness^1^. Extreme underdominance is an attractive and versatile tool for population control. First, it could be leveraged to create threshold-dependent, spatially-contained gene drives^2^ capable of replacing local populations. Such gene drives may be more socially acceptable than threshold-independent gene-drives to suppress populations since their spead can be more tightly controlled. Alternatively, only males could be released for a genetic biocontrol approach that mimics sterile insect technique. Several strategies for engineering underdominance have been described, including one- or two-locus toxin-antitoxin systems^3,4^, chromosomal translocations^5^, and RNAi-based negative genetic interactions^6^. Despite its theoretical utility in population control, engineering extreme underdominance has been difficult^1^.

Extreme underdominance essentially constitutes a speciation event, as it prevents successful reproduction and therefore genetic exchange between two populations. In nature, prezygotic and postzygotic incompatibilities maintain species barriers. Prezygotic incompatibilities prevent fertilization from taking place. These can include geographic separation or behavioral/anatomical differences between individuals in two populations that prevent sperm and egg from meeting. Postzygotic incompatibilities occur when genetic or cellular differences between the maternal and paternal gametes render the offspring inviable or infertile. The Dobzhansky-Muller Incompatibility (DMI) model^7,8^ asserts that postzygotic incompatibilities can arise via mutations that create a two-locus underdominance effect^9^. DMIs are considered a major driving force underlying natural speciation events. Understanding the molecular mechanisms resulting in hybrid incompatibilities between species is a central question for evolutionary biology and ecology.

We have recently described a versatile and effective method for engineering DMIs in the lab to direct synthetic speciation events. We name this method engineered genetic incompatibility (EGI). An EGI strain is homozygous for a lethal effector gene and the corresponding resistance allele(s). What separates EGI from described toxin/anti-toxin systems is that the lethal effector allele is haplosufficient, while the resistance allele is haploinsufficient. Any outcrossing of the EGI strain with wild-type generates inviable hybrids, as the resulting heterozygotes contain the lethal effector gene but only one copy of the haploinsufficient resistance allele (**Fig. 1a**). Unlike single locus, bi-allelic toxin-antitoxin systems^3^, the EGI genotype in principle incurs no fitness penalty, as 100% of the offspring between similarly engineered EGI parents remain viable. Our approach leverages sequence-programmable transcription activators (PTAs) to drive lethal over- or ectopic-expression of endogenous genes (**Fig. 1b/c**)^10^.

**Figure 1.**
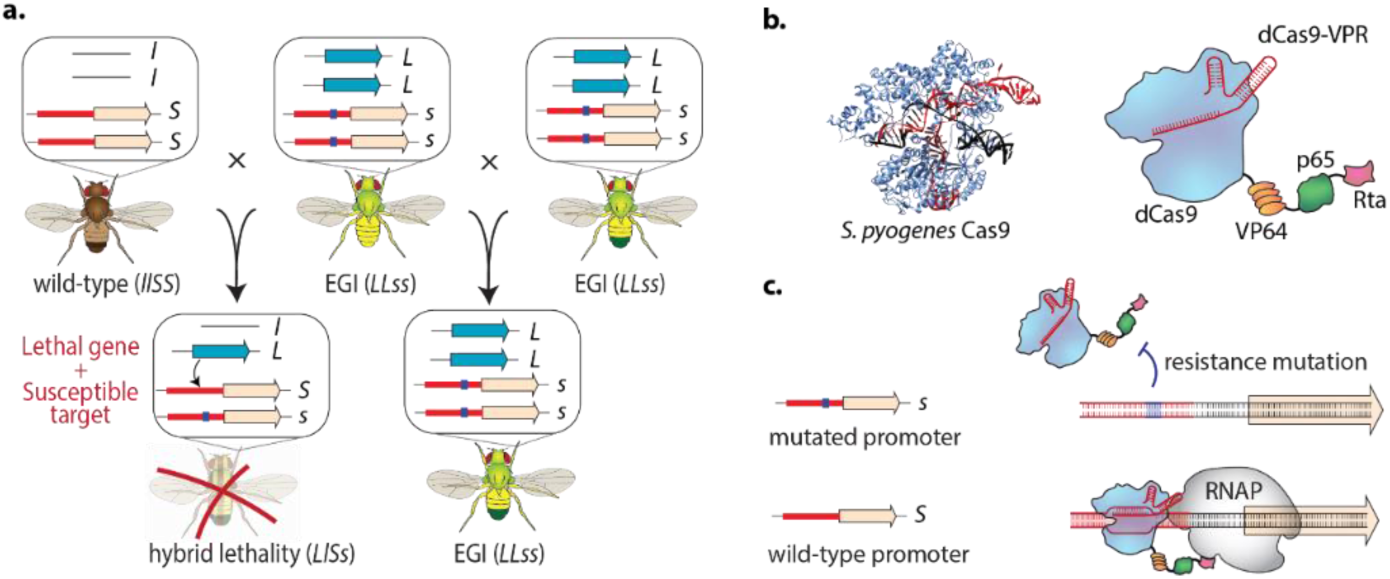
Design of Engineered Genetic Incompatibility (EGI). (**a**) Schematic diagram of required genotypes for EGI. *L*, dominant lethal gene; *l*, wild-type allele (null); *S*, dominant susceptible allele; *s*, recessive resistant allele. (**b**) X-ray crystal structure of *S. pyogenes* Cas9 (PDB ID: 6o0z, left) and diagram of dominant lethal gene product, dCas9-VPR. (**c**) Interaction of dCas9-VPR with resistant (top) or susceptible (bottom) alleles. RNAP, RNA polymerase.

Here we apply EGI to engineer extreme underdominance in the model insect, *Drosophila melanogaster*. We show that the strength and timing of hybrid lethality can be tuned based on genetic design. Further, we show that multiple mutually-incompatible ‘synthetic species’ can be created for the same target organism. This has important ramifications for the design of genetic biocontrol strategies that are robust in the face of genetic resistance.

### Lethal overexpression of endogenous genes

To drive lethal overexpression of endogenous genes, we use the dCas9-VPR, composed of a catalytically inactive Cas9 fused to three transcriptional activation domains (VP64, p65, and Rta)^11^ (**Fig. 1b**). This has been used previously to cause lethal gene activation in *D. melanogaster*^12^; however, we had to mitigate apparent off-target toxicity associated with strong dCas9-VPR expression in the absence of sgRNA^13^. Replacing the promoter driving dCas9-VPR with promoters from various developmental genes (*Pwg*, Pfoxo, Pbam*) or a truncated tubulin promoter (*Ptub*)^14^ allowed us to constrain dCas9-VPR expression sufficiently to allow generation of homozygous fly strains. We also created viable homozygous flies expressing the evolved dXCas9-VPR transactivator from the truncated tubulin promoter^15^.

Strains homozygous for dCas9-VPR constructs were mated to strains homozygous for sgRNAs targeting several genes important for development (*hh, hid, pyr, upd1, upd2, upd3, wg, vn*). The parental flies were removed from mating vials after five days and the number of offspring surviving to pupal and adult life-stages were counted after 15 days (**Fig. 2, Supplementary Data File 1**). Several crosses produced no surviving adult offspring in replicate experiments. Interestingly, we observed unique hybrid incompatibility phenotypes that depended on the combination of PTA and sgRNA used to drive over- or ectopic-expression. Six crosses (red shading, **Fig. 2**) yielded no pupae, indicating embryonic or larval lethality. The strongest early lethality was seen when *Ptub:dCas9-VPR* or *Pwg*:dCas9-VPR* drove expression of the developmental genes *pyramus, wingless*, and *unpaired*-1. Thirteen crosses (yellow shading, **Fig. 2**) produced a pupal-lethal phenotype. These include genotypes which are predominantly larval lethal with a small number of offspring surviving to form pupae (*e.g. Pfoxo:dCas9-VPR* with *wg-sgRNA*) as well as genotypes that give normal numbers of pupae forming, but no adults emerged (*e.g. Ptub:dCas9-VPR* with *upd3-sgRNA*). One of the crosses, *Pwg*:dCas9-VPR* X *upd3-sgRNA* (blue shading, **Fig. 2**) produced a small number of surviving adults that were visibly deformed and died within three days of emerging from pupae. Interestingly, we observed two crosses with the *Ptub:dXCas9-VPR* parent (green shading, **Fig. 2**) that showed strong sex-ratio biasing, with predominantly (95%, *upd1*) or exclusively (100%, *upd2*) male adult survivors. The same PTA crossed with *vn* exhibits a slight sex-ratio bias of 1.7:1 males:females (Supplementary Data File). We used these data to select a sub-set of putative target genes for constructing EGI flies, focusing on *pyr, wg*, and *hh* moving forward.

**Figure 2.**
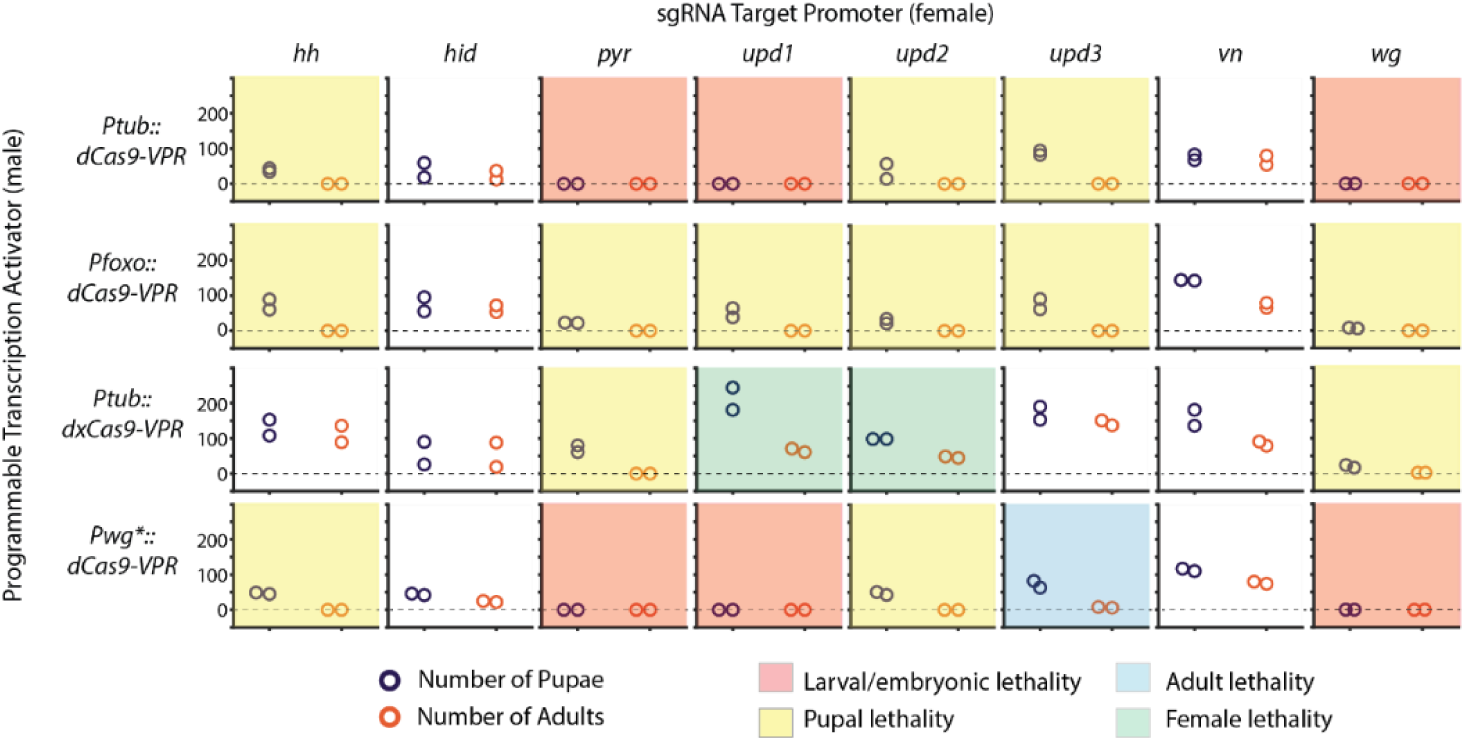
Empirical determination of targets for lethal over- or ectopic-expression. Results showing the number of progeny surviving to pupal or adult life-stages (purple, orange circles, respectively) for crosses between a paternal fly homozygous for a dCas9-VPR expression cassette (rows) and a maternal fly homozygous for sgRNA expression cassette (columns). Individual experiments are colored according to phenotype categories according to the key below.

### Constructing EGI strains

Mutations to the sgRNA-binding sequences of target promoters are necessary to prevent lethal over- or ectopic-expression in the EGI strain. These constitute the haploinsufficient resistance alleles in the EGI design (**Fig. 1a**,**c**). To generate viable promoter mutations, flies expressing germline active Cas9 nuclease^16^ were crossed to homozygous sgRNA-expressing strains^17^ or were directly microinjected with sgRNA expression constructs. Offspring were crossed to balancers and F2 flies were screened for the presence of mutations via Sanger sequencing. For each target promoter, we isolated mutant flies that were viable as homozygotes and without any apparent phenotype, suggesting that the mutations are benign and do not substantially interfere with required expression from these loci (**Fig. 3a, Supplementary Fig. S1**). It is noteworthy that we commonly recovered mutated promoters that had independent NHEJ events at each sgRNA target site, despite their close proximity. This is contrary to the belief that targeting proximal sequences is likely to result in complete excision of the intervening sequence in the event of NHEJ^18^.

**Figure 3.**
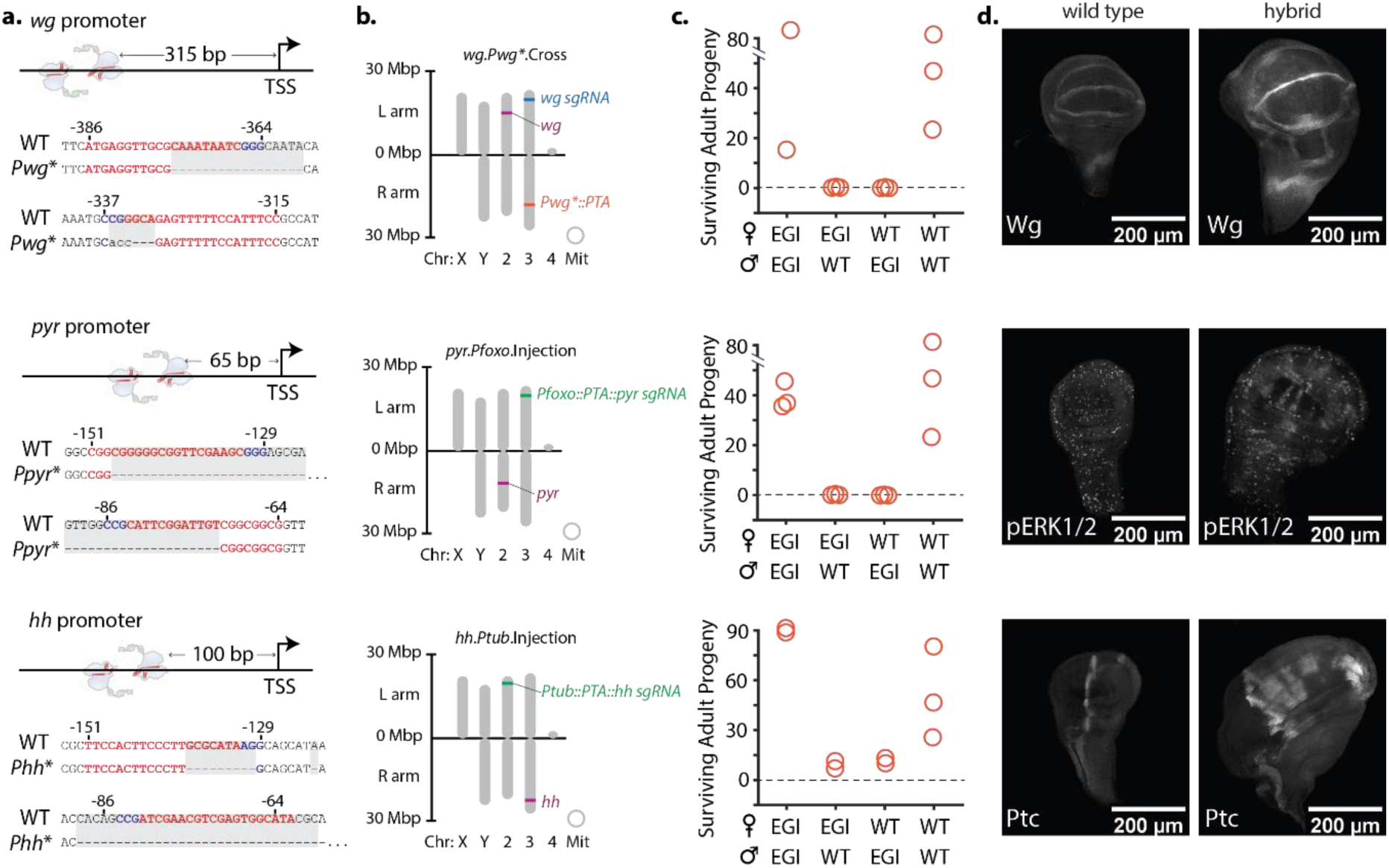
Genotype and hybrid incompatibility of select EGI strains. (**a**) Proximity of sgRNA binding sites to transcription start site (TSS) for EGI strains. Sequences of both sgRNA binding sites are shown below promoter illustration, with protospacers in red and protospacer adjacent motifs in blue. Sequences of the mutated promoters at the sgRNA binding loci are shown below with differences highlighted in grey shadow. (**b**) Chromosomal locations of genome alterations. (**c**) Hybrid incompatibility data showing number of progeny surviving to adulthood. Genotype of parental strains for each cross are given on the x-axis. (**d**) Immunohistochemical staining of wild-type (left) or hybrid (right) larva showing over- or ectopic-expression of targeted signalling pathways. Antibody binding targets are labelled in the bottom left corner of each image. For each panel, the *wg-, pyr-*, and *hh*-targeting EGI genotypes are shown from top to bottom.

We combined each of the required components to create a full EGI genotype via one of two approaches. In each, we needed to avoid passing through intermediate genotypes that contained an active PTA and a wild-type promoter sequence, as this would be lethal. The first method involved a total of 19 crosses between flies containing Cas9, PTA, or sgRNA expression constructs that had already been characterized in **Fig. 2** (**Fig. 3b top, Supplementary Figs. S2-S3**). The second method involved re-injecting embryos from homozygous promoter mutant strains with a single plasmid containing expression constructs for both the dCas9-VPR and the sgRNA (**Fig. 3b bottom**). The latter approach was more direct, requiring approximately half the number of crosses, but resulted in different chromosomal location for PTA expression compared to what was previously characterized (**Supplementary Figs. S4-S6**). Using these two methods, we produced a total of 12 unique EGI genotypes (**Supplementary Fig. S7**). We use a short-hand naming convention that describes the target gene (*wg, pyr, hh*), the promoter driving *dCas9-VPR* (*Pwg*, Pfoxo, Ptub, Pbam*), and the method used in strain construction (crossing, injection): for example *pyr.Pfoxo*.injection.

### Assessing Hybrid Incompatibility

Candidate EGI strains were crossed to wild-type (Oregon R and w1118) to assess mating compatibility. While w1118 was the ‘wild-type’ starting point for our EGI engineering efforts, male w1118 flies have a previously reported courtship phenotype^19^. Oregon R males lack this mating phenotype and reproduce more efficiently. We performed intra-specific matings (male and female from the same EGI genotype) and EGI × wild-type matings by combining three virgin females of one genotype with two males of another genotype. The number of pupae and adult progeny were counted after 15 days just as for the hybrid lethality screen described above. EGI strains that drove over- or ectopic-expression of *wingless* or *pyramus* both showed full incompatibility, with no hybrids surviving to adulthood (**Fig. 3c**). These represent engineered extreme underdominance. The EGI lines were healthy and fecund, with EGI × EGI crosses yielding numbers of offspring on par with wild-type × wild-type crosses. A third EGI strain targeting the *hedgehog* promoter showed a marked underdominant phenotype, but not as strong as the extremely underdominant *wg-* or *pyr-*EGI strains. Approximately 10-13% of hybrid offspring from *hh*-EGI crosses survived to adulthood. Of these surviving offspring, the females were all sterile, but the males were not, supporting a role of proper hedgehog expression in oogenesis^20,21^. That the initial *hh*-EGI strain was not as robust as *wg-* or *pyr-*EGI strains is not surprising. Activation of *hedgehog* produced a later-acting lethal phenotype compared to activation of *pyramus* and *wingless* in the PTA × sgRNA crosses (yielding pupal lethality instead of larval lethality). We believe that the weaker phenotype for *hh*-targeting guides in the EGI × wild-type hybrids (i.e. **Fig. 3**) versus the PTA × sgRNA crosses (i.e. **Fig. 2**) is the result of having only one sensitive (wild-type) promoter from which to drive lethal expression in the EGI × wild-type hybrids.

In order to confirm the mechanism of hybrid lethality, we performed immunohistochemistry on hybrid larva. We stained for target gene overexpression (Wingless) or activation of known downstream components in the relevant signalling pathways (p-ERK1*/2* and Patched (Ptc) for EGI targets *pyr* and *hh*, respectively). For our *wg*-targeting EGI line we observed overexpression, but no ectopic expression, in the wing imaginal disc as expected from our *Pwg*::dCas9-VPR* expression design, in which the PTA is itself driven by the mutated *wg* promoter (**Fig. 3d, top panel**). Interestingly, we did observe unique staining patterns in the brain, but are not sure if this is due to ectopic expression or just accumulation of the overproduced ligand (**Supplementary Fig. S8**). When we drive expression of *pyr* or *hh* with a *foxo* or short *tubulin* promoter, respectively, we observe clear evidence of ectopic expression in hybrid larva (**Fig. 3d, Supplementary Fig. S8**). For the *pyr*-targeting EGI line, we observed ectopic activation of pERK1/2 in clusters of cells throughout the wing imaginal disc, whereas pERK1/2 is normally activated in a speckled like pattern. For the *hh-*targeting EGI line, we observe Patched ectopic production only in the anterior compartment, which phenocopies previous experiments of *hh* overexpression in imaginal discs^22^.

### Mutual incompatibility between EGI strains with distinct genotypes

We predicted that our method of generating species-like barriers to sexual reproduction would allow us to engineer not just one, but many EGI genotypes that are all incompatible with wild-type and with each other. To test this, we performed a large all-by-all cross-compatibility experiment that included 12 EGI and 2 wild-type genotypes. Each cross was performed bi-directionally (female of strain A to male of strain B and vice versa). The orthogonality plot (**Fig. 4**) shows number of surviving adults from each cross. Crosses that are expected to produce viable offspring are present on the diagonal, with multiple “compatibility groups” defined by target-promoter mutations. Nine EGI strains were 100% incompatible with one or both wild-type lines. These include strains designed towards each of the developmental morphogen targets (*hh, pyr*, and *wg*). The high degree of symmetry across the diagonal shows that EGI produces bi-directional incompatibility, with the number of surviving offspring being similar if the EGI constructs were inherited maternally or paternally. While this is true when assessing the number of surviving adult progeny, we observed differences in timing of lethality for maternally versus paternally inherited EGI constructs (**Supplementary Data File 1, Supplementary Video File 1**).

**Figure 4.**
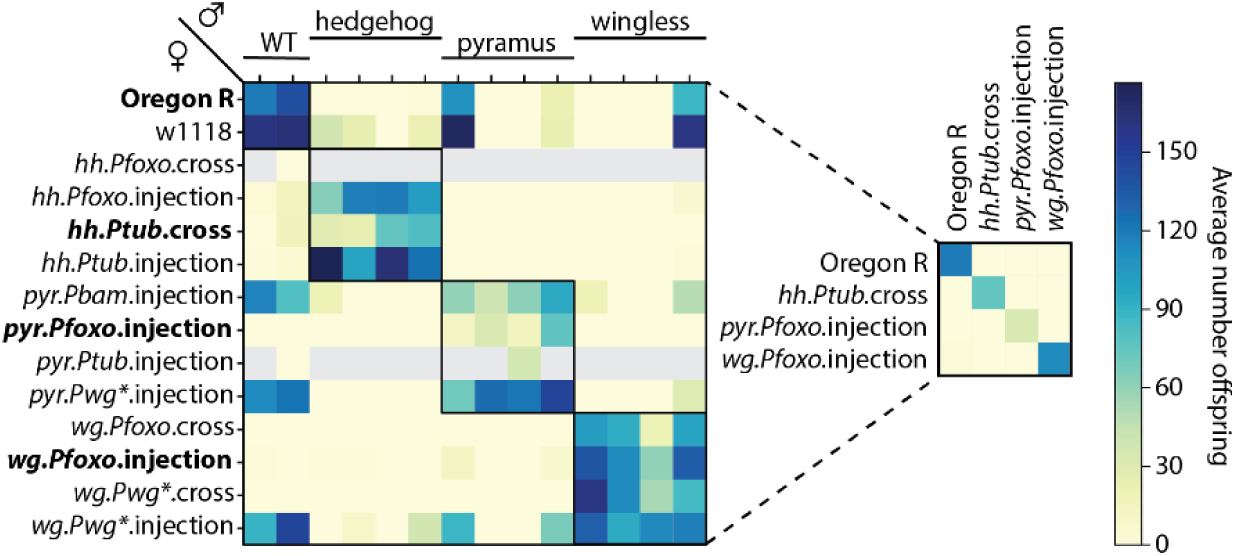
Engineering multiple orthogonal EGI strains. Mating compatibility between wild-type and 12 EGI genotypes, reported as the number of adult offspring 15 days after mating. Female (maternal) genotype is listed on the left axis with the naming convention [target.PTApromoter.construction-method], and male (paternal) genotypes are presented in the same order along the top axis. Predicted compatible strains are indicated with black-outline boxes across the diagonal. Grey boxes indicate crosses that were not measured for lack of virgin females for *hh.Pfoxo*.injection and *pyr.Ptub*.injection strains. Smaller grid at right highlights four mutually-compatible strains. Unless otherwise noted in **Supplementary Data File 1**, values represent mean of three independent replicates.

Nuanced differences in genetic design are important for the performance of EGI strains. For example, *wg.Pwg*\*.cross and *wg.Pwg*\*.injection have the same genetic components and are expected to work via identical mechanisms. The only difference is the chromosomal location of the PTA components. Despite their similarity, the former shows 100% incompatibility with wild-type, while the latter shows normal numbers of surviving offspring. This difference in performance is likely due to variable expression levels of the PTA construct, although this was not directly tested. Overall, these results show that we can engineer multiple EGI strains for a target organism. This has important implications in overcoming resistance to genetic population control, which is discussed in detail below.

## Discussion

Here we demonstrate the ability to rationally engineer species-like barriers to sexual reproduction in a multicellular organism. We employed the EGI approach that was recently described in yeast^10^. Our successful implementation in flies confirms that this is a broadly applicable strategy for engineering reproductive barriers. Engineered speciation has been previously described in *D. melanogaster* by Moreno, wherein a non-essential transcription factor, *glass*, was knocked out and *glass*-dependent lethal gene construct was introduced^23^. This approach also uses a similar topology to EGI; however, the resulting flies were blind in the absence of *glass* and this approach could not be scaled to make multiple incompatible strains. Our use of PTAs to drive lethal over- or ectopic-expression allows us to generate multiple EGI strains with no noticeable phenotypes aside from their hybrid incompatibility. Using a similar approach, Windbichler et al. developed PTAs capable of driving lethal overexpression of developmental morphogens in *D. melanogaster*, but were unable to generate complete EGI strains due to target selection and transgene toxicity^14^. We found that the ability create viable EGI lines requires empirical tweaking of genetic designs to ensure the required components are expressed at sufficiently high levels to drive lethal expression in hybrid offspring, but not so high to cause toxicity in the EGI organisms.

The ability to rationally design reproductive barriers opens up diverse opportunities for pest management and biocontrol of invasive species as well as genetic containment of novel proprietary genetically modified organisms^24^. Underdominance-based gene drives are threshold-dependent and allow for localized population replacement^25^. Several strategies for engineered underdominance exist^4–6^, but the EGI system is the first to produce 100% lethality of F1 heterozygotes. Compared to homing endonuclease gene drives, underdominance drives are more easily reversible and less likely to spread beyond the local target population^26^. Extreme underdominance gene drives are unique in their ability to spread genes/traits through a population that are unlinked to the drive allele. Since no hybrids between the biocontrol agent and wild-type organisms are viable, there is no opportunity for recombination events to break the linkage between the drive allele and other genes in the genome. Thus, EGI could be used to replace multi-locus traits in a target population.

Alternatively, EGI could be used as an alternative to Sterile Insect Technique^27^ by releasing only one sex. For example, released males would compete with wild males to mate with wild females. Any egg fertilized by an EGI male would fail to develop to adulthood. In applications such as this, our ability to tune the life-stage of hybrid lethality could have dramatic impact on the success of a biocontrol program. Late-acting pupal lethality would still allow for hybrid larva to compete for resources with wild-type larva. This is preferred for insects with ‘overcompensating density-dependence’ at larval stages^28–30^, where decreasing larval numbers increases the larval survival probability to the point where the total population actually grows instead of shrinks. On the contrary, embryonic-lethality could be preferred for agricultural pests whose larva cause extensive crop damage^31^.

There are three primary mechanisms by which the incompatibility provided by EGI could ‘break’: (i) transgene silencing of the dCas9-based PTA, (ii) early promoter conversion of the target locus in hybrid organisms, and (iii) underlying sequence diversity at target loci in wild populations that prevent PTA recognition. We have previously published an engineering solution to (i) that involves creating a positive selection for the PTA using endogenous essential genes^10^. Promoter conversion is unlikely to effect the success of biocontrol programs due to an inherent fitness defect in such escape mutants^32^. To address the underlying sequence diversity at target loci, population genetics studies should precede strain engineering to identify highly conserved targetable regions, which we have found to be present in populations of interest [Smanski, unpublished]. However, all of these resistance mechanisms can be mitigated using mutually incompatible EGI strains (*e.g.* **Fig. 4**). With just two orthogonal EGI strains (‘A’ and ‘B’), an iterative release of A-B-A-B-A-B… for biocontrol is expected to result in negatively correlated cross-resistance^33^ (**Supplementary Fig. S9**). Any surviving hybrids from mating events between EGI-A and wild-type would automatically inherit a susceptibility to EGI-B (because EGI-A and EGI-B are mutually incompatible). Thus these surviving ‘escapees’ would be sensitive to the next release of EGI-B, and this renewed sensitization would continue with each sequential release.

In summary, we demonstrate the ability to engineer species-like genetic incompatibilities in a multicellular organism. Our approach uses genetic tools that have been proven effective in many organisms, and our design is applicable to any sexually-reproducing species. We show that the EGI approach is robust to specific design implementations, with extreme underdominance possible with at least three distinct developmental morphogen targets. Further, we show that multiple ‘synthetic species’ can be engineered from a given target organism.

## Methods

### Plasmids

Plasmids expressing dCas9-VPR were constructed by Isothermal assembly^34^ combining NotI linearized pMBO2744 attP vector backbone with dCas9-VPR PCR amplified from pAct:dCas9-VPR (Addgene #78898)^35^ and SV40 terminator for pH-Stinger (BDSC, #1018) to generate pMM7-6-1. Isothermal assembly was used to clone 5’UTR and ∼1.5kb of promoter sequence into NotI linearized pMM7-6-1 (pMM7-6-2: Foxo promoter. pMM7-6-3: Tubulin promoter. pMM7-6-4: wingless promoter. pMM7-6-5: Bam promoter). Plasmids expressing dXCas9-VPR were constructed by introducing mutations into the dCas9 region predicted to improve activity^36^ to generate pMM7-9-3 which also has a NotI linearization site used for cloning promoter and 5’UTR sequences.

Plasmids expressing sgRNAs were generated by cloning annealed oligos into p{CFD4-3xP3::DsRed} (Addgene #86864).

Plasmids expressing both sgRNAs and dCas9-VPR were constructed by Isothermal assembly combining KpnI linearized dCas9-VPR plasmids (pMM7-6-2 through pMM7-6-5) with sgRNAs amplified from genomic DNA from *Drosophila melanogaster* stocks that are available from BDSC (pyr sgRNA: 67537. Hh sgRNA: 67560. Upd1: 67555. Wg: 67545). See **Supplementary Table S2** for plasmid descriptions.

### Drosophila stocks

Experimental crosses were performed at 25°C and 12 hour days. Existing Cas9 and sgRNA strains were obtained from the Bloomington Drosophila Stock Center. All transgenic flies were generated via ΦC31 mediated integration targeted to attP landing sites. Embryo microinjections were performed by BestGene Inc (Chino Hills, Ca). See **Supplementary Table S1** for descriptions of fly strains.

### Drosophila rearing conditions

All drosophila strains were grown on Bloomington Formulation Nutri-Fly media containing 4% v/v 1M propionic acid (pH 4.3). Additional dry yeast crumbs were added to vials during EGI strain generation matings. No additional yeast was used in any mating compatibility tests. The flies were housed at 25° C with 12hr day/night cycles. Drosophila strains used in the all-by-all cross were moved to 18° C overnight to aid in virgin female collection the following day.

### Mating compatibility tests

Genetic compatibility was assayed between parental stock homozygous for the PTA or sgRNA expression cassette (i.e. PTA-sgRNA testing) as well as between final EGI genotypes and wild-type (i.e. EGI testing). Test crosses were performed by crossing sexually-mature adult males to sexually-mature virgin females homozygous for their respective genotype at a ratio of 3:3 (PTA-sgRNA testing) or 2:3 (EGI testing). The adults were removed from the vials after 5 days and the offspring were counted after 15 days. Filled and empty pupal cases were counted towards the pupae total and adult males and females were counted towards the adult count. Independent mating compatibility tests were performed in duplicate (PTA-sgRNA testing) or triplicate (EGI testing). In all-by-all EGI compatibility test (**Fig. 4**), the data for the *pyr.Pfoxo*.injection self-cross was performed independently from the other crosses in that dataset.

### Immunohistochemistry

Late 3rd instar larvae were dissected in cold PBS, and fixed with 4% formaldehyde (Electron Microscopy Science, RT-15714) overnight at 4°C. Tissues were washed and permeabilized with PBS-TritonX-100 (0.1%) before staining with appropriate antibodies. Tissues for fluorescence microscopy were mounted with 80% Glycerol in PBS (0.1% TritonX-100). Images were captured using the Zeiss LSM710. Confocal Z-stacks were processed in FIJI (ImageJ).

### Antibodies and staining reagents

Drosophila-Patched, apa1 (Developmental Studies Hybridoma Bank (DSHB)) (1:50); Drosophila-Wingless, 4D4 (DSHB) (1:50); Drosophila-Armadillo, N2-7A1 (DSHB) (1:50); Phospho-MAPK (ERK1/2), #4370 (Cell Signaling Technologies) (1:100). AlexaFluor 568 and 647 (Invitrogen) conjugated secondary antibodies were used as necessary at (1:500) dilution. Tissues were counterstained with DAPI (Millipore Sigma, #D9542) (1 µg/ml).

## Acknowledgements

MJS is supported in part by the Defense Advanced Research Projects Agency (grant number D17AP00028). The views, opinions, and/or findings contained in this article are those of the authors and should not be interpreted as representing the official views or policies, either expressed or implied, of the Defense Advanced Research Projects Agency or the Department of Defense.

## Author contributions

MM and MJS conceived this study. MM, NF, AH, AU, AJP, MBO, and MJS designed experiments. MM, NF, AU, AH, SD, and NM performed experiments. MM, NF, and MJS wrote the manuscript.

## Supplementary Information for

## This PDF file includes

Supplementary Figs. S1 to S9

Supplementary Tables S1 to S4

Legend for Supplementary Data File 1

Legend for Supplementary Video File 1

References

## Other Supplementary Materials for this manuscript include the following

Supplementary Data File 1

Supplementary Video File 1

**Supplementary Figure S1.**
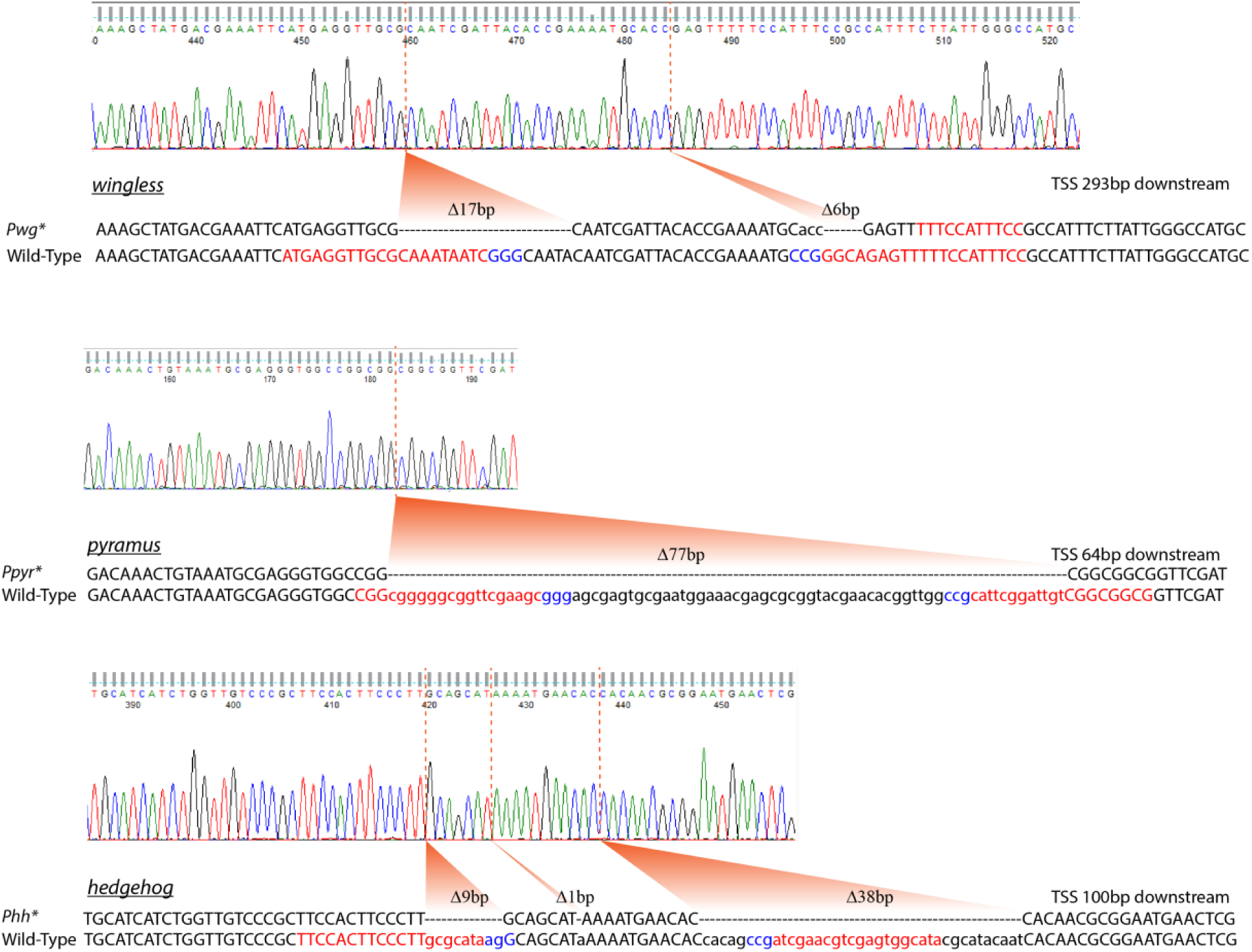
Promoter Mutations. Promoter mutant sequencing traces and alignments to wild-type promoters. Targeted protospacers indicated in red and PAMs indicated in blue.

**Supplementary Figure S2.**
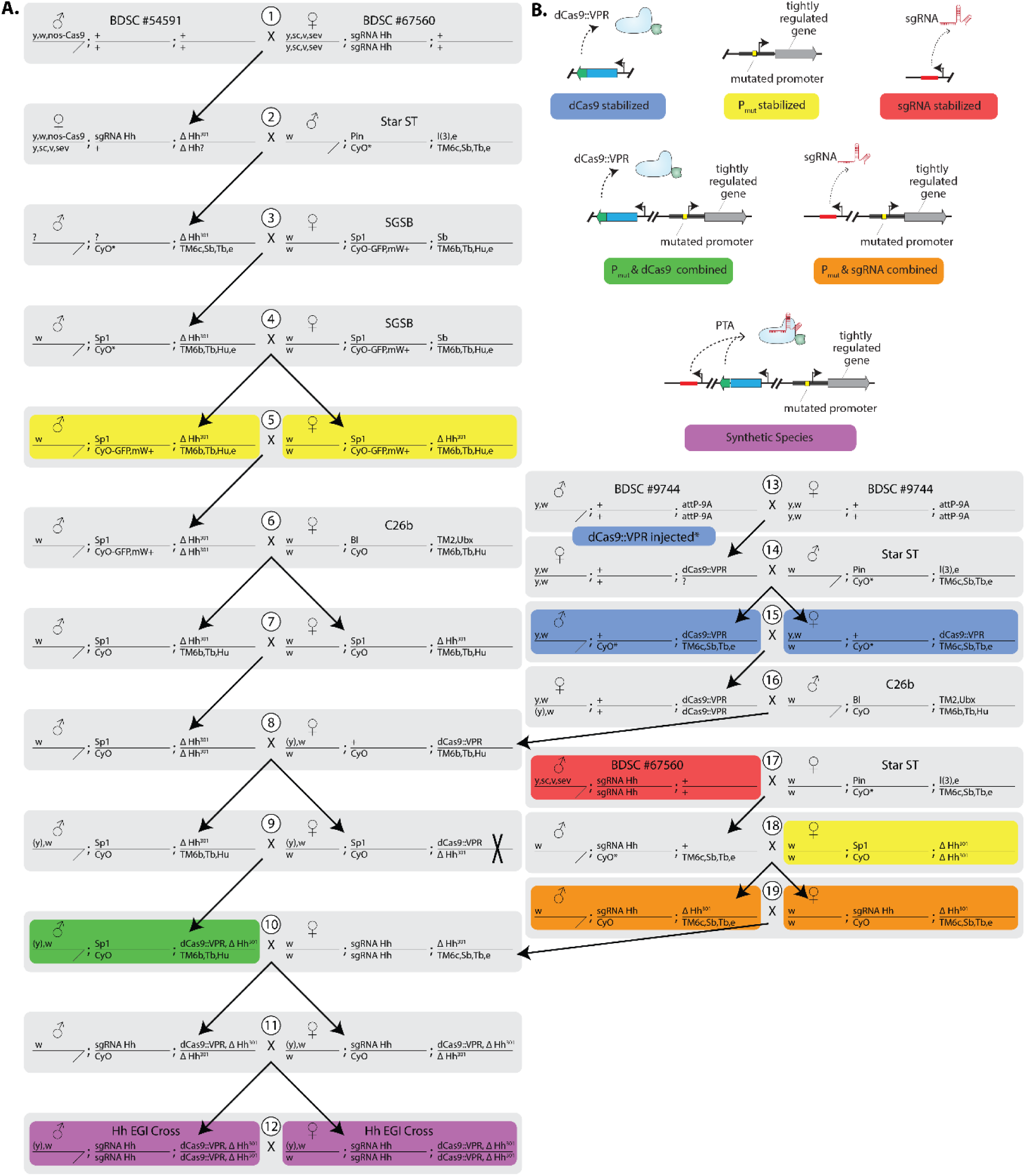
Crossing strategy to produce hh-EGI flies. (A.) Genotypes and sex of flies involved in crosses required to bring together EGI components. Crosses are indexed with numbered white circles. ‘X’ designates a recombination event required in the female parent of cross #9. The female from cross #18 resulted from cross #7 (not cross #5). Embryos from cross #13 were injected with promoter::dCas9::VPR constructs and ΦC31 integrase. Question mark denotes a chromosome genotype that was not verified. (B.) Illustrated and color-coded genotypes of key intermediates. BDSC #54591, BDSC #67560, and BDSC #9744 were purchased from the Bloomington Drosophila Stock Center. Star ST, SGSB, and C26b are balancer strains.

**Supplementary Figure S3.**
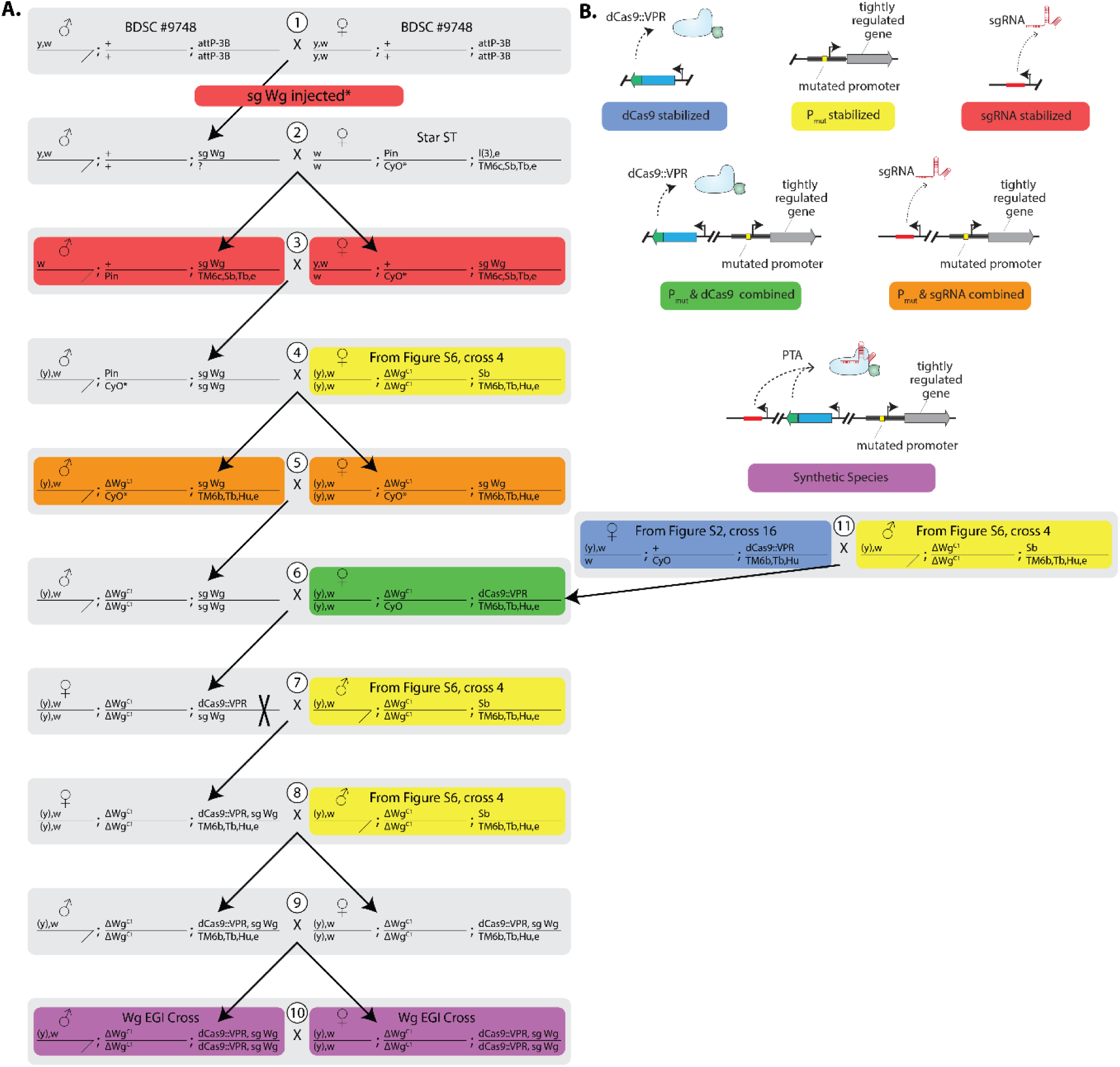
Crossing strategy to produce wg-EGI flies. (A.) Genotypes and sex of flies involved in crosses required to bring together EGI components. Crosses are indexed with numbered white circles. ‘X’ designates a recombination event required in the female parent of cross #7. Embryos from cross #1 were injected with a sgRNA-wg construct and ΦC31 integrase. Question mark denotes a chromosome genotype that was not verified. The males in crosses #7, 8, and 11 and the female in cross #4 are offspring from Suppl. Fig. S6, cross #4. The female in cross #11 is offspring from Suppl. Fig. S2, cross #16. (B.) Illustrated and color-coded genotypes of key intermediates. BDSC #9748 was purchased from the Bloomington Drosophila Stock Center. Star ST is a balancer strain.

**Supplementary Figure S4.**
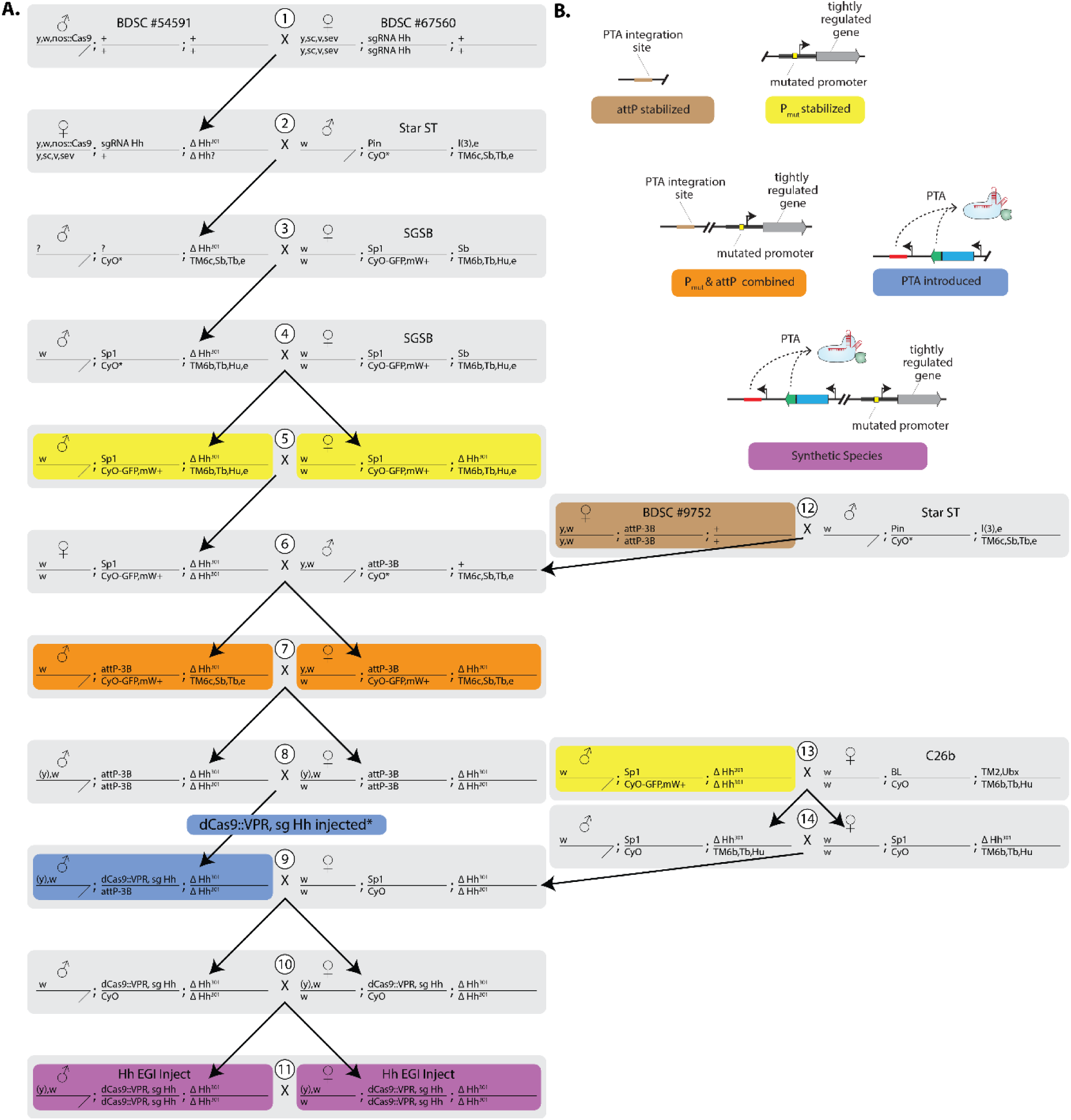
Reinjection strategy to produce hh-EGI flies. (A.) Genotypes and sex of flies involved in crosses required to bring together EGI components. Crosses are indexed with numbered white circles. Embryos from cross #8 were injected with promoter::dCas9::VPR + sgRNA-hh constructs and ΦC31 integrase. Question mark denotes a chromosome genotype that was not verified. (B.) Illustrated and color-coded genotypes of key intermediates. BDSC #54591, BDSC #67560, and BDSC #9752 were purchased from the Bloomington Drosophila Stock Center. Star ST, SGSB, and C26b are balancer strains.

**Supplementary Figure S5.**
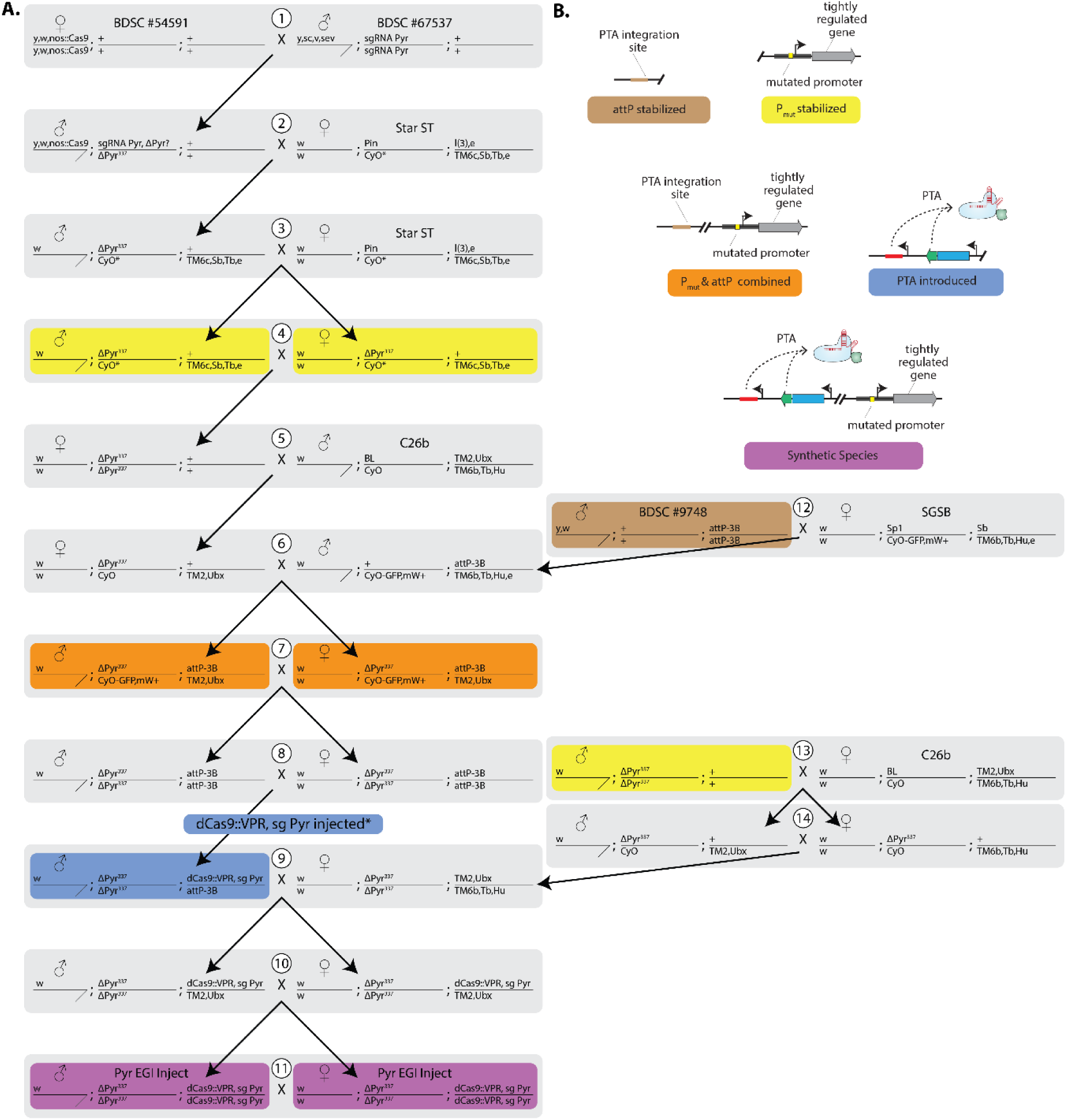
Reinjection strategy to produce pyr-EGI flies. (A.) Genotypes and sex of flies involved in crosses required to bring together EGI components. Crosses are indexed with numbered white circles. Embryos from cross #8 were injected with promoter::dCas9::VPR + sgRNA-pyr constructs and ΦC31 integrase. Question mark denotes a chromosome genotype that was not verified. (B.) Illustrated and color-coded genotypes of key intermediates. BDSC #54591, BDSC #67537, and BDSC #9748 were purchased from the Bloomington Drosophila Stock Center. Star ST, SGSB, and C26b are balancer strains.

**Supplementary Figure S6.**
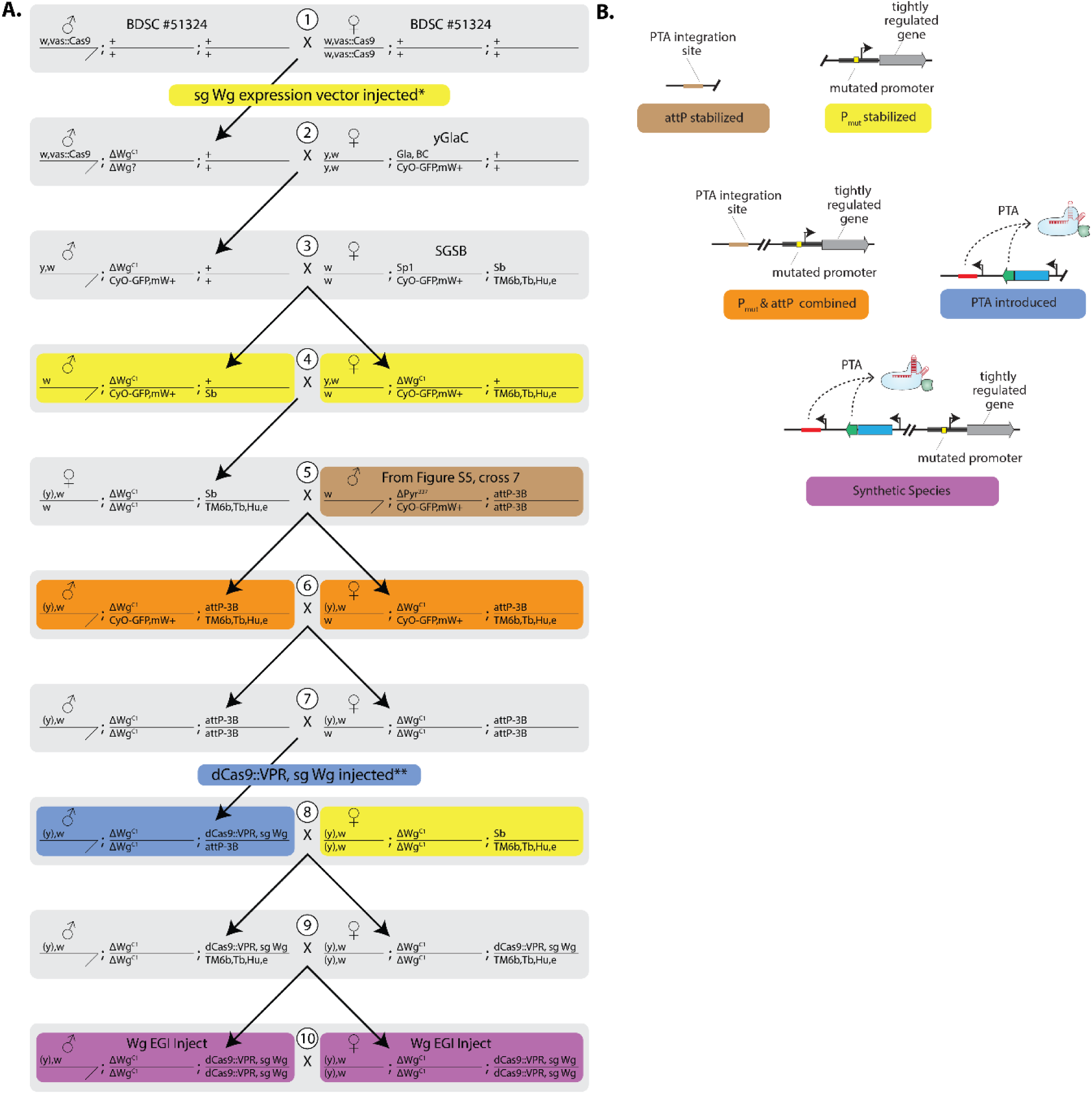
Reinjection strategy to produce wg-EGI flies. (A.) Genotypes and sex of flies involved in crosses required to bring together EGI components. Crosses are indexed with numbered white circles. Embryos from cross #1 were injected with the sgRNA-wg expression construct. Embryos from cross #7 were injected with promoter::dCas9::VPR + sgRNA-wg constructs and ΦC31 integrase. Question mark denotes a chromosome genotype that was not verified. The male in cross #5 is offspring from Suppl. Fig. S5, cross #7. (B.) Illustrated and color-coded genotypes of key intermediates. BDSC #51324 was purchased from the Bloomington Drosophila Stock Center. yGlac and SGSB are balancer strains.

**Supplementary Figure S7:**
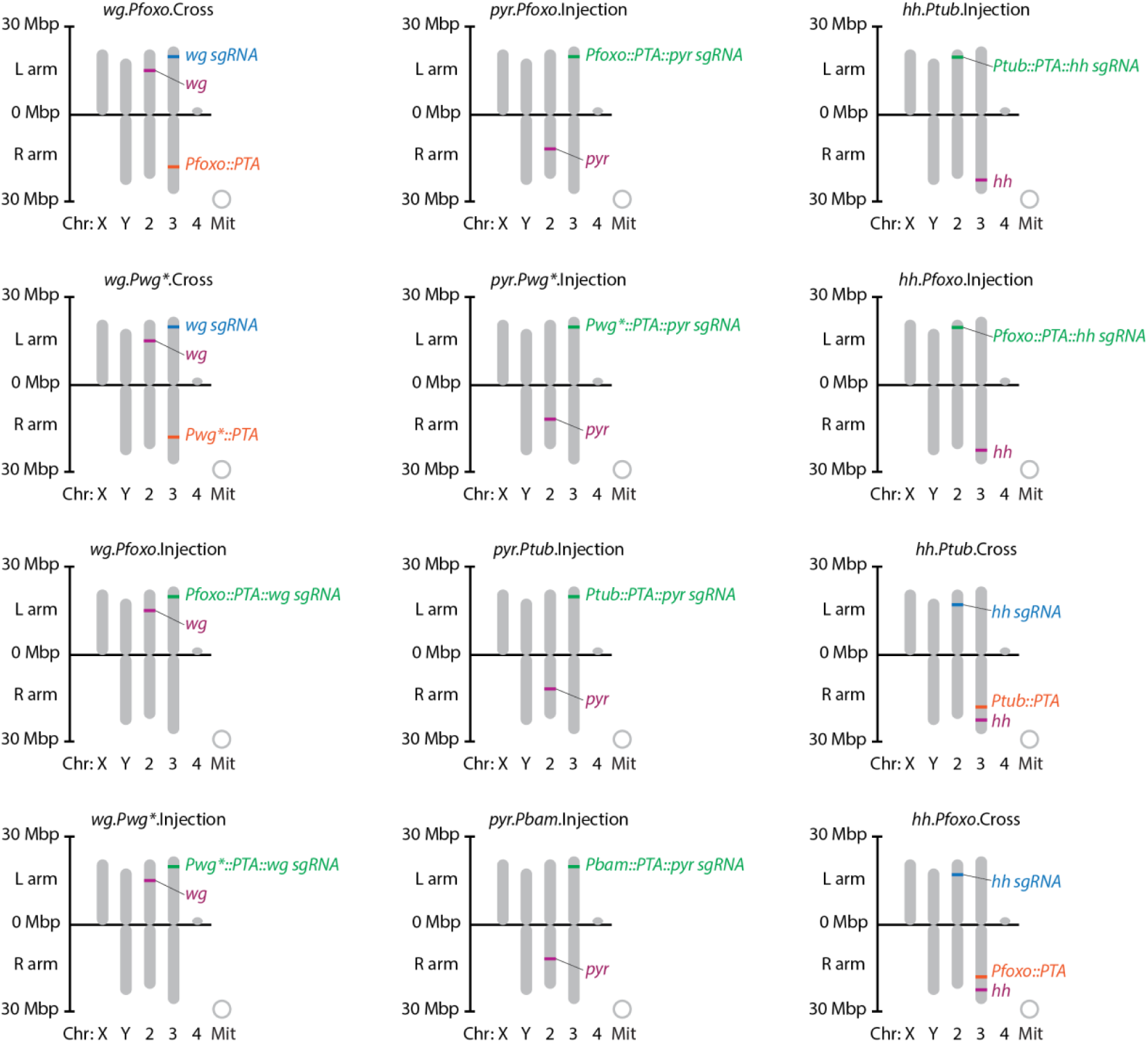
Chromosomal maps of all EGI strains reported in this work.

**Supplementary Fig. S8:**
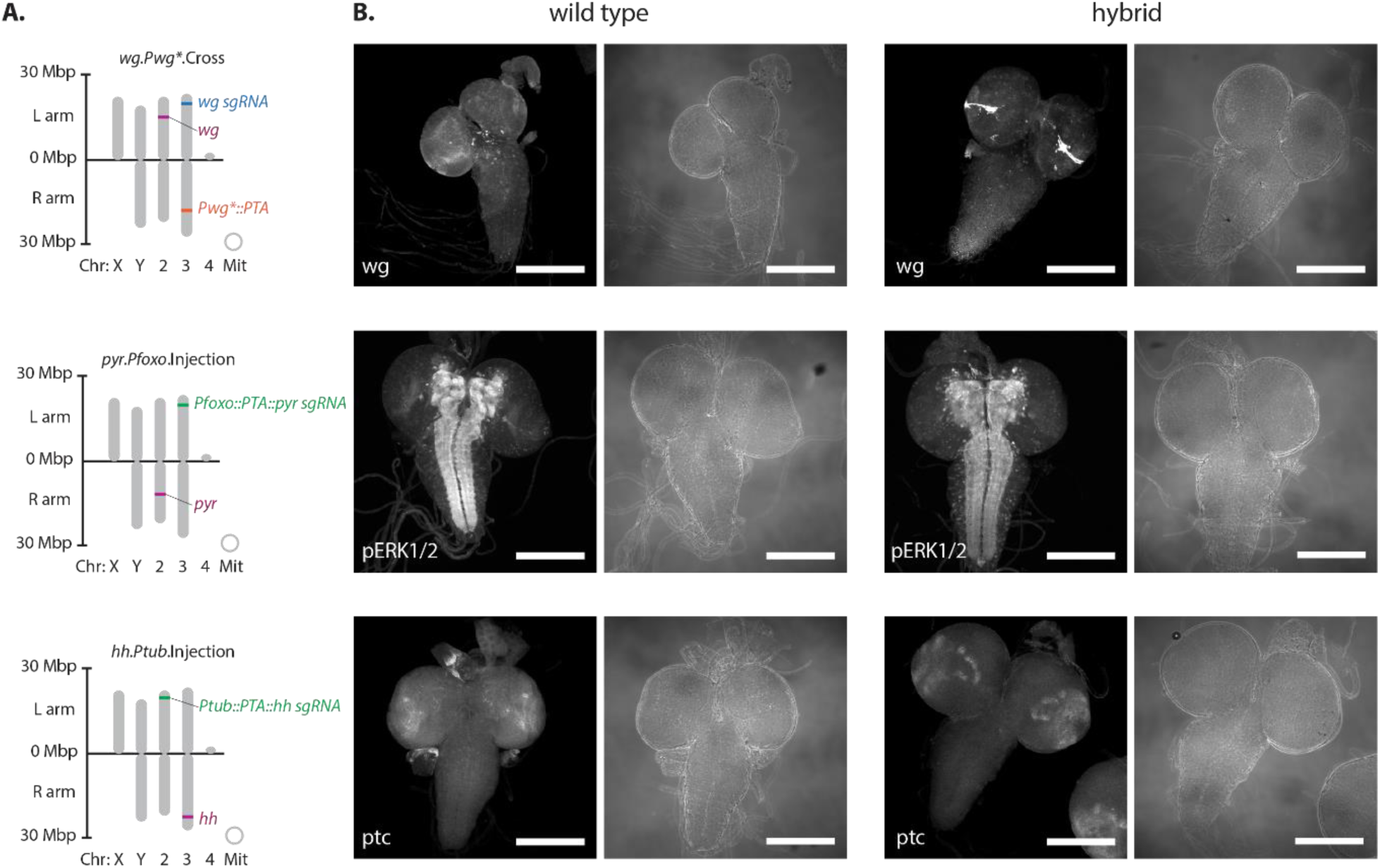
(A.) Chromosomal locations of genome alterations for EGI strains whose hybrid offspring were analysed by immunohistochemistry. EGI strains illustrated here correspond to the ones used in Fig. 3. (B.) Immunofluorescence staining of 3rd instar larval brains from wild-type (left) or hybrid (right) showing over- or ectopic-expression of targeted signalling pathways. Grayscale images show antibody staining for proteins encoded by lethal overexpression target (*wingless*, top) or downstream signalling pathway components (p-*ERK1/2*, middle and *patched*, bottom). Corresponding brightfield images of the brains to the right. Scale bar = 200 µm.

**Supplementary Figure S9.**
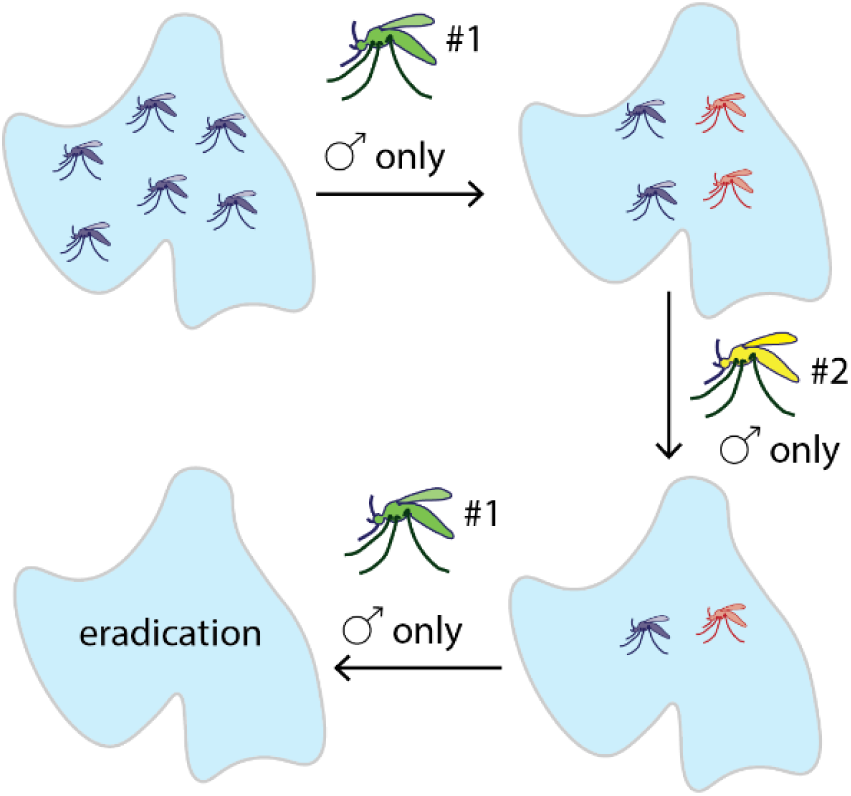
Release scheme for negatively correlating cross-resistance. Purple denotes wild-type pests, green and yellow denote mutually-incompatible EGI strains, for which only males would be released. Orange denotes resistant ‘escapees’, which inherit half of their genome from the previously released biocontrol EGI strain.

**Supplementary Table S1:**
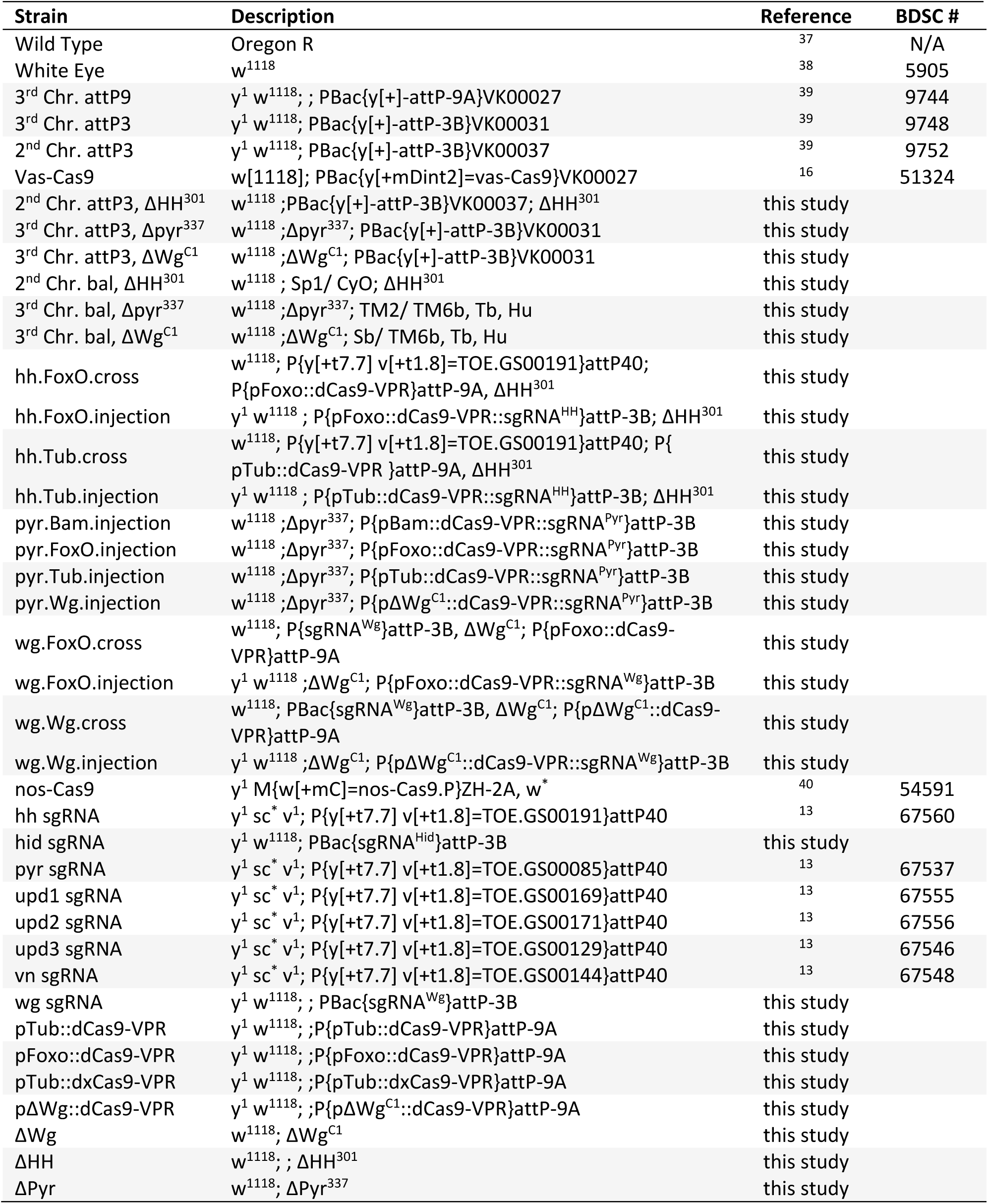
*D. melanogaster* strains used in this study.

| Strain | Description | Reference | BDSC # |
| --- | --- | --- | --- |
| Wild Type | Oregon R | 37 | N/A |
| White Eye | w<sup>1118</sup> | 38 | 5905 |
| 3<sup>rd</sup> Chr. attP9 | y<sup>1</sup> w<sup>1118</sup>; ; PBac{y[+]-attP-9A}VK00027 | 39 | 9744 |
| 3<sup>rd</sup> Chr. attP3 | y<sup>1</sup> w<sup>1118</sup>; PBac{y[+]-attP-3B}VK00031 | 39 | 9748 |
| 2<sup>nd</sup> Chr. attP3 | y<sup>1</sup> w<sup>1118</sup>; PBac{y[+]-attP-3B}VK00037 | 39 | 9752 |
| Vas-Cas9 | w[1118]; PBac{y[+mDint2]=vas-Cas9}VK00027 | 16 | 51324 |
| 2<sup>nd</sup> Chr. attP3, ΔHH<sup>301</sup> | w<sup>1118</sup>; PBac{y[+]-attP-3B}VK00037; ΔHH<sup>301</sup> | this study |  |
| 3<sup>rd</sup> Chr. attP3, Δpyr<sup>337</sup> | w<sup>1118</sup>; Δpyr<sup>337</sup>; PBac{y[+]-attP-3B}VK00031 | this study |  |
| 3<sup>rd</sup> Chr. attP3, ΔWg<sup>C1</sup> | w<sup>1118</sup>; ΔWg<sup>C1</sup>; PBac{y[+]-attP-3B}VK00031 | this study |  |
| 2<sup>nd</sup> Chr. bal, ΔHH<sup>301</sup> | w<sup>1118</sup>; Sp1/ CyO; ΔHH<sup>301</sup> | this study |  |
| 3<sup>rd</sup> Chr. bal, Δpyr<sup>337</sup> | w<sup>1118</sup>; Δpyr<sup>337</sup>; TM2/ TM6b, Tb, Hu | this study |  |
| 3<sup>rd</sup> Chr. bal, ΔWg<sup>C1</sup> | w<sup>1118</sup>; ΔWg<sup>C1</sup>; Sb/ TM6b, Tb, Hu | this study |  |
| hh.FoxO.cross | w<sup>1118</sup>; P{y[+t7.7] v[+t1.8]=TOE.GS00191}attP40; P{pFoxo::dCas9-VPR}attP-9A, ΔHH<sup>301</sup> | this study |  |
| hh.FoxO.injection | y<sup>1</sup> w<sup>1118</sup>; P{pFoxo::dCas9-VPR::sgRNA<sup>HH</sup>}attP-3B; ΔHH<sup>301</sup> | this study |  |
| hh.Tub.cross | w<sup>1118</sup>; P{y[+t7.7] v[+t1.8]=TOE.GS00191}attP40; P{pTub::dCas9-VPR }attP-9A, ΔHH<sup>301</sup> | this study |  |
| hh.Tub.injection | y<sup>1</sup> w<sup>1118</sup>; P{pTub::dCas9-VPR::sgRNA<sup>HH</sup>}attP-3B; ΔHH<sup>301</sup> | this study |  |
| pyr.Bam.injection | w<sup>1118</sup>; Δpyr<sup>337</sup>; P{pBam::dCas9-VPR::sgRNA<sup>Pyr</sup>}attP-3B | this study |  |
| pyr.FoxO.injection | w<sup>1118</sup>; Δpyr<sup>337</sup>; P{pFoxo::dCas9-VPR::sgRNA<sup>Pyr</sup>}attP-3B | this study |  |
| pyr.Tub.injection | w<sup>1118</sup>; Δpyr<sup>337</sup>; P{pTub::dCas9-VPR::sgRNA<sup>Pyr</sup>}attP-3B | this study |  |
| pyr.Wg.injection | w<sup>1118</sup>; Δpyr<sup>337</sup>; P{pΔWg<sup>C1</sup>::dCas9-VPR::sgRNA<sup>Pyr</sup>}attP-3B | this study |  |
| wg.FoxO.cross | w<sup>1118</sup>; P{sgRNA<sup>Wg</sup>}attP-3B, ΔWg<sup>C1</sup>; P{pFoxo::dCas9-VPR}attP-9A | this study |  |
| wg.FoxO.injection | y<sup>1</sup> w<sup>1118</sup>; ΔWg<sup>C1</sup>; P{pFoxo::dCas9-VPR::sgRNA<sup>Wg</sup>}attP-3B | this study |  |
| wg.Wg.cross | w<sup>1118</sup>; PBac{sgRNA<sup>Wg</sup>}attP-3B, ΔWg<sup>C1</sup>; P{pΔWg<sup>C1</sup>::dCas9-VPR}attP-9A | this study |  |
| wg.Wg.injection | y<sup>1</sup> w<sup>1118</sup>; ΔWg<sup>C1</sup>; P{pΔWg<sup>C1</sup>::dCas9-VPR::sgRNA<sup>Wg</sup>}attP-3B | this study |  |
| nos-Cas9 | y<sup>1</sup> M{w[+mC]=nos-Cas9.P}ZH-2A, w<sup>*</sup> | 40 | 54591 |
| hh sgRNA | y<sup>1</sup> sc<sup>*</sup> v<sup>1</sup>; P{y[+t7.7] v[+t1.8]=TOE.GS00191}attP40 | 13 | 67560 |
| hid sgRNA | y<sup>1</sup> w<sup>1118</sup>; PBac{sgRNA<sup>Hid</sup>}attP-3B | this study |  |
| pyr sgRNA | y<sup>1</sup> sc<sup>*</sup> v<sup>1</sup>; P{y[+t7.7] v[+t1.8]=TOE.GS00085}attP40 | 13 | 67537 |
| upd1 sgRNA | y<sup>1</sup> sc<sup>*</sup> v<sup>1</sup>; P{y[+t7.7] v[+t1.8]=TOE.GS00169}attP40 | 13 | 67555 |
| upd2 sgRNA | y<sup>1</sup> sc<sup>*</sup> v<sup>1</sup>; P{y[+t7.7] v[+t1.8]=TOE.GS00171}attP40 | 13 | 67556 |
| upd3 sgRNA | y<sup>1</sup> sc<sup>*</sup> v<sup>1</sup>; P{y[+t7.7] v[+t1.8]=TOE.GS00129}attP40 | 13 | 67546 |
| vn sgRNA | y<sup>1</sup> sc<sup>*</sup> v<sup>1</sup>; P{y[+t7.7] v[+t1.8]=TOE.GS00144}attP40 | 13 | 67548 |
| wg sgRNA | y<sup>1</sup> w<sup>1118</sup>; ; PBac{sgRNA<sup>Wg</sup>}attP-3B | this study |  |
| pTub::dCas9-VPR | y<sup>1</sup> w<sup>1118</sup>; ; P{pTub::dCas9-VPR}attP-9A | this study |  |
| pFoxo::dCas9-VPR | y<sup>1</sup> w<sup>1118</sup>; ; P{pFoxo::dCas9-VPR}attP-9A | this study |  |
| pTub::dxCas9-VPR | y<sup>1</sup> w<sup>1118</sup>; ; P{pTub::dxCas9-VPR}attP-9A | this study |  |
| pΔWg::dCas9-VPR | y<sup>1</sup> w<sup>1118</sup>; ; P{pΔWg<sup>C1</sup>::dCas9-VPR}attP-9A | this study |  |
| ΔWg | w<sup>1118</sup>; ΔWg<sup>C1</sup> | this study |  |
| ΔHH | w<sup>1118</sup>; ; ΔHH<sup>301</sup> | this study |  |
| ΔPyr | w<sup>1118</sup>; ΔPyr<sup>337</sup> | this study |  |

**Supplementary Table S2:**
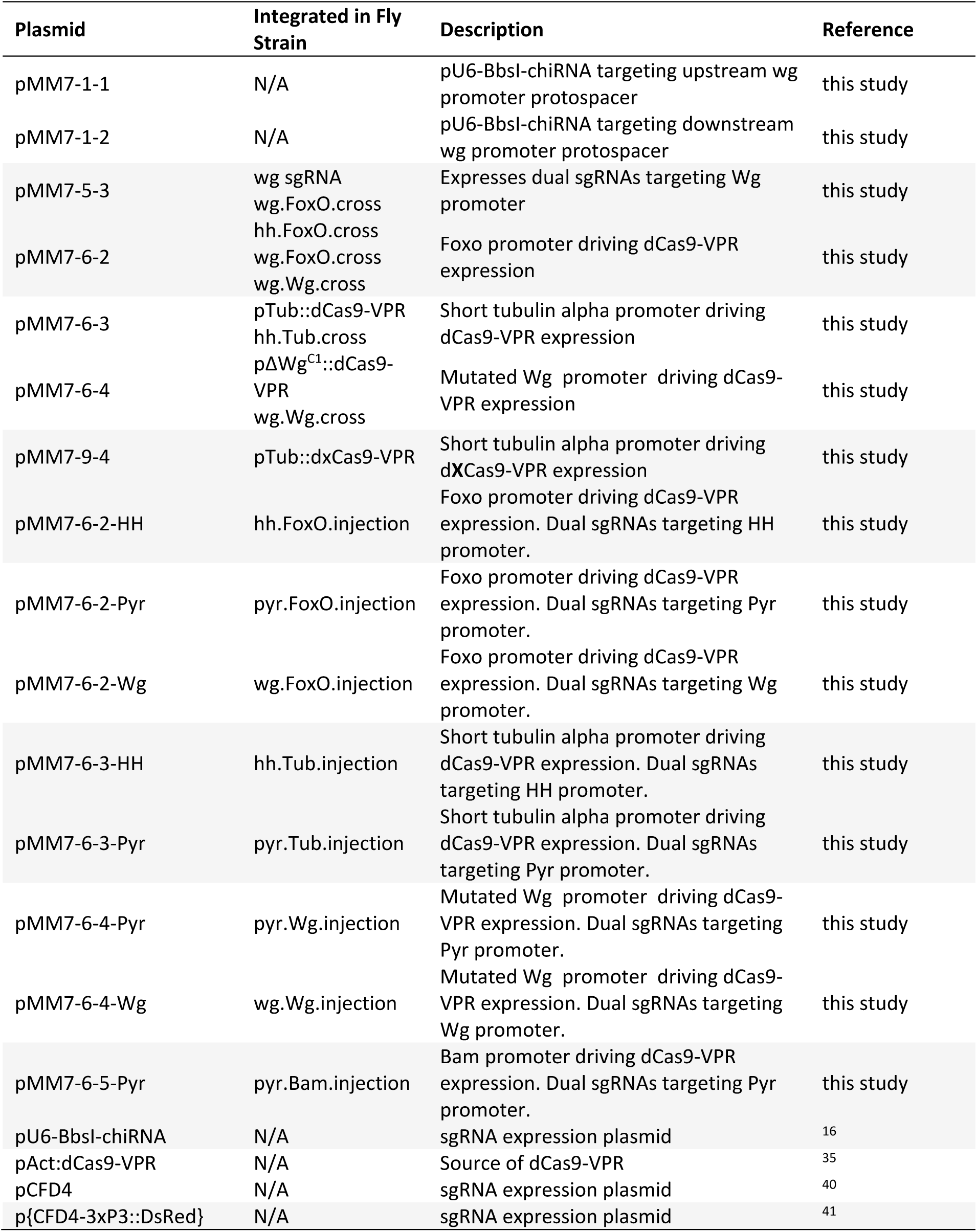
Plasmids used in this study.

**Supplementary Table S3:**
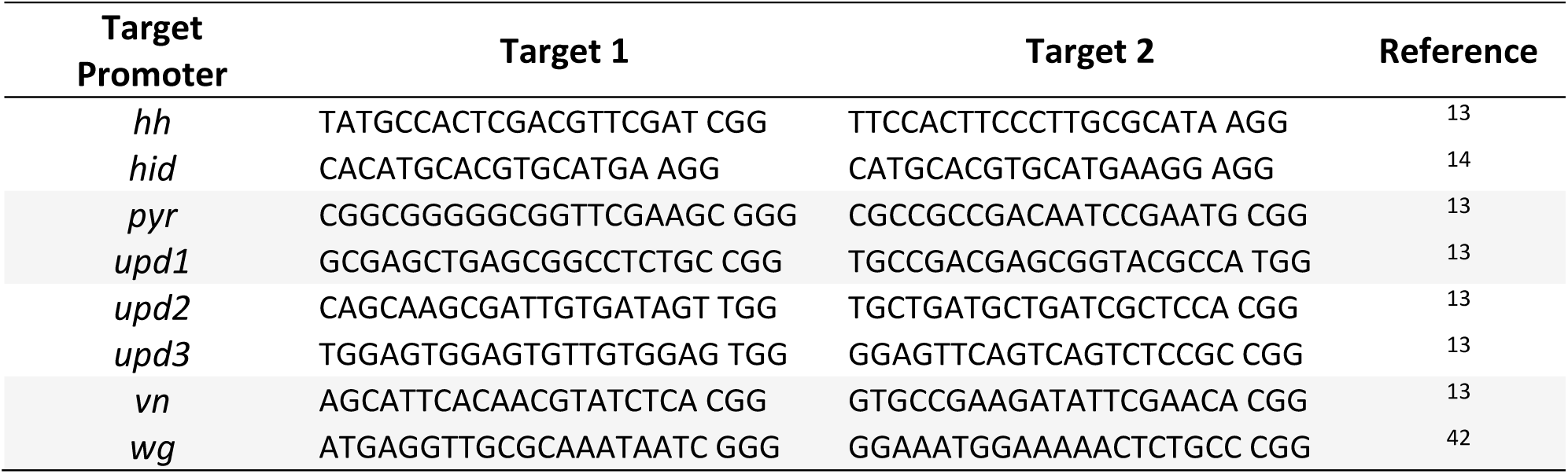
Promoter Target Sequences.

**Supplementary Table S4:**
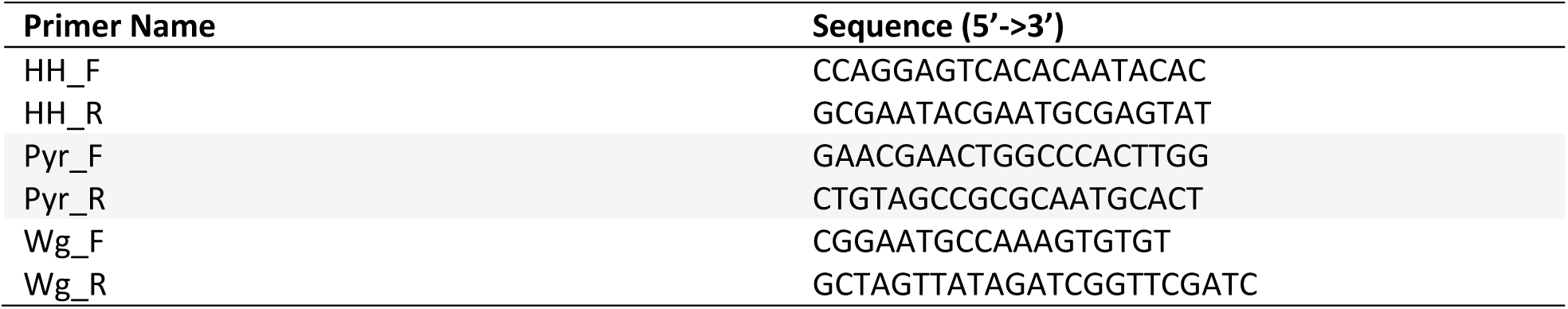
Promoter Sequencing Primers.

## Supplementary Data File (Attached as FlyEGI_SDF.xlsx)

The Supplementary Data File is an excel (.xlsx) file containing all of the numerical results from fly mating experiments described in this manuscript. There are three sheets, corresponding to data graphically reported in Figures2-4. Below is a description of the types of data contained in each sheet (bold) and column (italics).

### Sheet 1: PTA-sgRNA testing (figure 2)

*Column A. dCas9 Activator:* Description of the protein-only component of the PTA (i.e. dCas9 with no sgRNA) that is homozygous in the male adult involved in the cross

*ColumnB. gRNA target gene:* Description of the sgRNA construct that is homozygous in the female adult involved in the cross.

*Columns C,H. Pupae:* Counts of the total number of pupae from each cross, as determined by counting pupal casings. Data are reported separately for two biological replicates.

*Columns D,I. Males:* Counts of the total number of adult males present 15 days after crosses were set up. Data are reported separately for two biological replicates.

*Columns E,J. Females:* Counts of the total number of adult females present 15 days after crosses were set up. Data are reported separately for two biological replicates.

*Columns F,K. Stuck:* Counts of the adults that became stuck in the fly medium for which sex was not determined, 15 days after crosses were set up. Data are reported separately for two biological replicates.

*Columns G,L. Total Adults:* Sum of Males, Females and Stuck adults from previous columns. Data are reported separately for two biological replicates.

*Column M. Observed Lethality:* Assigned phenotype based on hybrid survival data.

### Sheet 2: EGI x WT testing (figure 3)

*Column A. Paternal genotype:* Strain name of the male adult involved in the cross.

*Column B. Maternal genotype:* Strain name of the female adult involved in the cross.

*Columns C,F,I. Pupae:* Counts of the total number of pupae from each cross, as determined by counting pupal casings. Data are reported separately for three biological replicates.

*Columns D,G,J. Males:* Counts of the total number of adult males present 15 days after crosses were set up. Data are reported separately for three biological replicates.

*Columns E,H,K. Females:* Counts of the total number of adult females present 15 days after crosses were set up. Data are reported separately for three biological replicates.

### Sheet 3: EGI testing (figure 4)

*Column A. Paternal genotype:* Strain name of the male adult involved in the cross.

*Column B. Maternal genotype:* Strain name of the female adult involved in the cross.

*Columns C,F,I. Pupae:* Counts of the total number of pupae from each cross, as determined by counting pupal casings. Data are reported separately for three biological replicates.

*Columns D,G,J. Males:* Counts of the total number of adult males present 15 days after crosses were set up. Data are reported separately for three biological replicates.

*Columns E,H,K. Females:* Counts of the total number of adult females present 15 days after crosses were set up. Data are reported separately for three biological replicates.

### Supplementary Video File (Attached as FlyEGI_SVF.pptx)

PowerPoint file containing an embedded time-lapse video of representative mating from **Fig. 3**. EGI flies are *pyr.Pfoxo*.Injection genotype. Images were taken on a Canon EOS Rebel T5 Digital SLR equipped with a Satechi remote shutter. Image files were compiled into movie (with shake correction) using Adobe Premier Pro 2019. For the purpose of the video, lights were left on for all 24 hours, 15 days of the experiment. During experiments reported in the paper, normal light-dark cycles were used, as described above.

**Supplementary References**

## References

(1) Davis, S.; Bax, N.; Grewe, P. (2001) Engineered Underdominance Allows Efficient and Economical Introgression of Traits into Pest Populations. J. Theor. Biol. 212 (1), 83–98. https://doi.org/10.1006/jtbi.2001.2357.

(2) Leftwich, P. T.; Edgington, M. P.; Harvey-Samuel, T.; Carabajal Paladino, L. Z.; Norman, V. C.; Alphey, L. Recent Advances in Threshold-Dependent Gene Drives for Mosquitoes. Biochemical Society Transactions. 2018. https://doi.org/10.1042/BST20180076.

(3) Sinkins, S. P.; Gould, F. (2006) Gene Drive Systems for Insect Disease Vectors. Nat. Rev. Genet. 7 (6), 427–435. https://doi.org/10.1038/nrg1870.

(4) Akbari, O. S.; Matzen, K. D.; Marshall, J. M.; Huang, H.; Ward, C. M.; Hay, B. A. (2013) A Synthetic Gene Drive System for Local, Reversible Modification and Suppression of Insect Populations. Curr. Biol. 23 (8), 671–677. https://doi.org/10.1016/j.cub.2013.02.059.

(5) Buchman, A. B.; Ivy, T.; Marshall, J. M.; Akbari, O. S.; Hay, B. A. (2018) Engineered Reciprocal Chromosome Translocations Drive High Threshold, Reversible Population Replacement in Drosophila. ACS Synth. Biol. 7 (5), 1359–1370. https://doi.org/10.1021/acssynbio.7b00451.

(6) Guy Reeves, R.; Bryk, J.; Altrock, P. M.; Denton, J. A.; Reed, F. A. (2014) First Steps towards Underdominant Genetic Transformation of Insect Populations. PLoS One. https://doi.org/10.1371/journal.pone.0097557.

(7) Dobzhansky, T. (1936) Studies on Hybrid Sterility II: Localization of Sterility Factors in Drosophila Pseudoobscura Hybrids. Genetics 21, 113–135.

(8) Muller, H. J. Isolating Mechanisms, Evolution and Temperature. Biol. Symp. 1942, p 71–125 ST–Isolating mechanisms, evolution and t.

(9) Orr, H. A.; Turelli, M. (2001) The Evolution of Postzygotic Isolation: AccumulatingDobzhansky-Muller Incompatibilities. Evolution (N. Y). 55 (6), 1085–1094.

(10) Maselko, M.; Heinsch, S. C.; Chacón, J. M.; Harcombe, W. R.; Smanski, M. J. (2017) Engineering Species-like Barriers to Sexual Reproduction. Nat. Commun. 8 (1). https://doi.org/10.1038/s41467-017-01007-3.

(11) Chavez, A.; Scheiman, J.; Vora, S.; Pruitt, B. W.; Tuttle, M.; P R Iyer, E.; Lin, S.; Kiani, S.; Guzman, C. D.; Wiegand, D. J.; et al. (2015) Highly Efficient Cas9-Mediated Transcriptional Programming. Nat. Methods. https://doi.org/10.1038/nmeth.3312.

(12) Lin, S.; Ewen-Campen, B.; Ni, X.; Housden, B. E.; Perrimon, N. (2015) In Vivo Transcriptional Activation Using CRISPR/Cas9 in Drosophila. Genetics 201 (2), 433–442. https://doi.org/10.1534/genetics.115.181065.

(13) Ewen-Campen, B.; Yang-Zhou, D.; Fernandes, V. R.; González, D. P.; Liu, L.-P.; Tao, R.; Ren, X.; Sun, J.; Hu, Y.; Zirin, J.; et al. (2017) Optimized Strategy for in Vivo Cas9-Activation in *Drosophila*. Proc. Natl. Acad. Sci. 201707635. https://doi.org/10.1073/pnas.1707635114.

(14) Waters, A. J.; Capriotti, P.; Gaboriau, D.; Papathanos, P. A.; Windbichler, N.; Alexander, S.; Building, F.; Kensington, S. (2018) Rationally-Engineered Reproductive Barriers Using CRISPR & CRISPRa: An Evaluation of the Synthetic Species Concept in Drosophila Melanogaster. Sci. Transl. Med. https://doi.org/10.1101/259010.

(15) Hu, J. H.; Miller, S. M.; Geurts, M. H.; Tang, W.; Chen, L.; Sun, N.; Zeina, C. M.; Gao, X.; Rees, H. A.; Lin, Z.; et al. (2018) Evolved Cas9 Variants with Broad PAM Compatibility and High DNA Specificity. Nature. https://doi.org/10.1038/nature26155.

(16) Gratz, S. J.; Cummings, A. M.; Nguyen, J. N.; Hamm, D. C.; Donohue, L. K.; Harrison, M. M.; Wildonger, J.; O’connor-Giles, K. M. Genome Engineering of Drosophila with the CRISPR RNA-Guided Cas9 Nuclease. Genetics. August 2013, pp 1029–1035. https://doi.org/10.1534/genetics.113.152710.

(17) Ewen-Campen, B.; Yang-Zhou, D.; Fernandes, V. R.; González, D. P.; Liu, L. P.; Tao, R.; Ren, X.; Sun, J.; Hu, Y.; Zirin, J.; et al. (2017) Optimized Strategy for in Vivo Cas9-Activation in Drosophila. Proc. Natl. Acad. Sci. U. S. A. https://doi.org/10.1073/pnas.1707635114.

(18) Champer, J.; Liu, J.; Oh, S. Y.; Reeves, R.; Luthra, A.; Oakes, N.; Clark, A. G.; Messer, P. W. (2018) Reducing Resistance Allele Formation in CRISPR Gene Drive. Proc. Natl. Acad. Sci. U. S. A. https://doi.org/10.1073/pnas.1720354115.

(19) Krstic, D.; Boll, W.; Noll, M. (2013) Influence of the White Locus on the Courtship Behavior of Drosophila Males. PLoS One. https://doi.org/10.1371/journal.pone.0077904.

(20) Forbes, A. J.; Lin, H.; Ingham, P. W.; Spradling, A. C. (1996) Hedgehog Is Required for the Proliferation and Specification of Ovarian Somatic Cells Prior to Egg Chamber Formation in Drosophila. Development.

(21) Vied, C.; Horabin, J. I. (2001) The Sex Determination Master Switch, Sex-Lethal, Responds to Hedgehog Signaling in the Drosophila Germline. Development.

(22) Alexandre, C.; Jacinto, A.; Ingham, P. W. (1996) Transcriptional Activation of Hedgehog Target Genes in Drosophila Is Mediated Directly by the Cubitus Interruptus Protein, a Member of the GLI Family of Zinc Finger DNA-Binding Proteins. Genes Dev. https://doi.org/10.1101/gad.10.16.2003.

(23) Moreno, E. (2012) Design and Construction of “Synthetic Species.” PLoS One 7 (7), e39054. https://doi.org/10.1371/journal.pone.0039054.

(24) Clark, M.; Maselko, M. (2020) Transgene Biocontainment Strategies for Molecular Farming. Front. Plant Sci. 11 (March), 1–11. https://doi.org/10.3389/fpls.2020.00210.

(25) Magori, K.; Gould, F. (2006) Genetically Engineered Underdominance for Manipulation of Pest Populations: A Deterministic Model. 2620 (April), 2613–2620. https://doi.org/10.1534/genetics.105.051789.

(26) Champer, J.; Buchman, A.; Akbari, O. S. Cheating Evolution: Engineering Gene Drives to Manipulate the Fate of Wild Populations. Nature Reviews Genetics. 2016, pp 146–159. https://doi.org/10.1038/nrg.2015.34.

(27) Dyck, V. A.; Hendrichs, J.; Robinson, A. S. Sterile Insect Technique: Principles and Practice in Area-Wide Integrated Pest Management. https://doi.org/10.5860/CHOICE.43-5894.

(28) Alphey, N.; Bonsall, M. B. (2014) Interplay of Population Genetics and Dynamics in the Genetic Control of Mosquitoes. J. R. Soc. Interface. https://doi.org/10.1098/rsif.2013.1071.

(29) Legros, M.; Lloyd, A. L.; Huang, Y.; Gould, F. (2009) Density-Dependent Intraspecific Competition in the Larval Stage of *Aedes Aegypti* (Diptera: Culicidae): Revisiting the Current Paradigm. J. Med. Entomol. 46 (3), 409–419. https://doi.org/10.1603/033.046.0301.

(30) Walsh, R. K.; Facchinelli, L.; Ramsey, J. M.; Bond, J. G.; Gould, F. (2011) Assessing the Impact of Density Dependence in Field Populations of Aedes Aegypti. J. Vector Ecol. https://doi.org/10.1111/j.1948-7134.2011.00170.x.

(31) Asplen, M. K.; Anfora, G.; Biondi, A.; Choi, D. S.; Chu, D.; Daane, K. M.; Gibert, P.; Gutierrez, A. P.; Hoelmer, K. A.; Hutchison, W. D.; et al. (2015) Invasion Biology of Spotted Wing Drosophila (Drosophila Suzukii): A Global Perspective and Future Priorities. J. Pest Sci. (2004). 88 (3), 469–494. https://doi.org/10.1007/s10340-015-0681-z.

(32) Maselko, M.; Heinsch, S.; Das, S.; Smanski, M. J. (2018) Genetic Incompatibility Combined with Female-Lethality Is Effective and Robust in Simulations of Aedes Aegypti Population Control. Bioarxiv http://dx.doi.org/10.1101/316406.

(33) Chapman, R. B.; Penman, D. R. Negatively Correlated Cross-Resistance to a Synthetic Pyrethroid in Organo-Phosphorus-Resistant Tetranychus Urticae [11]. Nature. 1979. https://doi.org/10.1038/281298a0.

(34) Gibson, D. G.; Young, L.; Chuang, R.-Y.; Venter, J. C.; Hutchison, C. A.; Smith, H. O. (2009) Enzymatic Assembly of DNA Molecules up to Several Hundred Kilobases. Nat. Methods 6, 343–345. https://doi.org/10.1038/nmeth.1318.

(35) Chavez, A.; Tuttle, M.; Pruitt, B. W.; Ewen-campen, B.; Chari, R.; Ter-ovanesyan, D.; Haque, S. J.; Cecchi, R. J.; Kowal, E. J. K.; Buchthal, J.; et al. (2016) Comparison of Cas9 Activators in Multiple Species. Nat. Methods No. May, 1–7. https://doi.org/10.1038/nmeth.3871.

(36) Richardson, C. D.; Ray, G. J.; DeWitt, M. A.; Curie, G. L.; Corn, J. E. (2016) Enhancing Homology-Directed Genome Editing by Catalytically Active and Inactive CRISPR-Cas9 Using Asymmetric Donor DNA. Nat. Biotechnol. https://doi.org/10.1038/nbt.3481.

(37) Lindsley, D. L.; Grell, E. H.; Bridges, C. B. (Calvin B. Genetic Variations of Drosophila Melanogaster.

(38) Ryder, E.; Hartl, D. L.; Wu, C. -tin. (2004) The DrosDel Collection: A Set of P-Element Insertions for Generating Custom Chromosomal Aberrations in Drosophila Melanogaster. Genetics 167 (2), 797–813. https://doi.org/10.1534/genetics.104.026658.

(39) Venken, K. J. T.; He, Y.; Hoskins, R. A.; Bellen, H. J. (2006) P[Acman]: A BAC Transgenic Platform for Targeted Insertion of Large DNA Fragments in D. Melanogaster. Science 314 (5806), 1747–1751. https://doi.org/10.1126/science.1134426.

(40) Port, F.; Chen, H. M.; Lee, T.; Bullock, S. L. (2014) Optimized CRISPR/Cas Tools for Efficient Germline and Somatic Genome Engineering in Drosophila. Proc. Natl. Acad. Sci. U. S. A. 111 (29), E2967–76. https://doi.org/10.1073/pnas.1405500111.

(41) Addgene: p{CFD4-3xP3::DsRed}.

(42) Lin, S.; Ewen-Campen, B.; Ni, X.; Housden, B. E.; Perrimon, N. (2015) In Vivo Transcriptional Activation Using CRISPR/Cas9 in Drosophila. Genetics 201 (2), 433–442. https://doi.org/10.1534/genetics.115.181065.

